# Human DNA replication initiation sites are specified epigenetically by oxidation of 5-methyl-deoxycytidine

**DOI:** 10.1101/2024.09.06.611651

**Authors:** Torsten Krude, Jiaming Bi, Rachel Doran, Rebecca A. Jones, James C Smith

## Abstract

DNA replication initiates at tens of thousands of sites on the human genome during each S phase. However, no consensus DNA sequence has been found that specifies the locations of these replication origins. Here, we investigate modifications of human genomic DNA by density equilibrium centrifugation and DNA sequencing. We identified short discrete sites with increased density during quiescence and G1 phase that overlap with DNA replication origins before their activation in S phase. The increased density is due to the oxidation of 5-methyl-deoxycytidines by TET enzymes at GC-rich domains. Reversible inhibition of de novo methylation and of subsequent oxidation of deoxycytidines results in a reversible inhibition of DNA replication and of cell proliferation. Our findings suggest a mechanism for the epigenetic specification and semiconservative inheritance of DNA replication origin sites in human cells that also provides a stable integral DNA replication licence to support once-per-cell cycle control of origin activation.

## Introduction

The replication of large eukaryotic genomes requires remarkable parallel processing. DNA replication forks are assembled and activated at large numbers of sites on linear chromosomes. It can be estimated, based on genome size and the speeds of DNA replication forks, that several tens of thousands of DNA replication start sites, or replication origins, are used in each human cell cycle (Hu and Stillman, 2023). The locations of these origins, and the mechanism by which they are specified in the human genome, are controversial topics.

Human DNA replication start sites appear to be organised hierarchically. Genome-wide DNA replication fork directionality analyses such as sequencing of Okazaki fragments (OK-seq) and leading strand 3’ ends (TrAEL-seq), or optical replication mapping of single-molecules (ORM), support the existence of broad initiation zones several tens of kilobases in size (Petryk et al., 2016; Kara et al., 2021; Wang et al., 2021; Wu et al., 2023). Within these large zones, DNA replication can initiate at several sites, often with relatively low efficiency.

Discrete and narrowly-defined initiation sites, however, have been mapped in human cells by several independent approaches. Short nascent strand sequencing (SNS-seq) is based on the isolation and sequencing of RNA-capped small nascent single-stranded DNA strands in replicating cells (Martin et al., 2011; Besnard et al., 2012; Picard et al., 2014; Akerman et al., 2020; Guilbaud et al., 2022). Initiation site sequencing (Ini-seq) involves labelling nascent DNA immediately after the initiation of DNA replication in a human cell-free system (Krude, 2000), followed by the isolation and sequencing of labelled double-stranded DNA fragments either by immunoprecipitation (Ini-seq version 1, (Langley et al., 2016)) or by density substitution and gradient centrifugation (Ini-seq version 2, (Guilbaud et al., 2022)). Both SNS-seq and Ini-seq have identified several tens of thousands of discrete replication initiation sites of only a few kilobases or less in the human genome, and there is generally good concordance between the locations of the discrete initiation sites identified by the different methods. The relationship between broad initiation zones and discrete initiation sites is unresolved and a subject of debate (Hu and Stillman, 2023). However, discrete and highly efficient initiation sites, as defined by Ini-seq (version 2), often demarcate the borders of initiation zones and additional, less efficient initiation sites are present within (Guilbaud et al., 2022; Murat et al., 2022).

It is not known how these discrete replication initiation sites in human cells are specified. In the budding yeast *S. cerevisiae*, early genetic experiments established that short DNA sequence elements called autonomous replicating sequences (ARS elements) are sufficient to attract the DNA replication initiation machinery and allow replication of any colinear DNA (Stinchcomb et al., 1979; Hsiao and Carbon, 1979; Hu and Stillman, 2023). Importantly, such sequence-specific ARS elements have not been found in mammalian cells. Genome-wide correlation studies, however, agree that active DNA replication origins are generally GC-rich and their genomic locations often correlate with CpG islands, with so-called ‘origin G-rich Repeated Element’ (OGRE) motifs, G quadruplexes or epigenetic chromatin marks that include open chromatin and certain histone modifications (Martin et al., 2011; Besnard et al., 2012; Valton et al., 2014; Picard et al., 2014; Cayrou et al., 2015; Langley et al., 2016; Prorok et al., 2019; Akerman et al., 2020; Guilbaud et al., 2022). G quadruplexes are formed by stacked guanosine tetrads and generate complex secondary structures involving looped single-stranded DNA domains on the G-rich strand and on the opposite C-rich strand. They have been shown to demarcate some DNA replication start sites and to facilitate de novo initiation when transplanted to different regions of the genome (Valton et al., 2014; Prioleau, 2017; Prorok et al., 2019; Poulet-Benedetti et al., 2023). DNA in CpG islands is often methylated symmetrically on both strands at the 5’ positions of cytosines, and methylated CpG islands are implicated in the regulation of gene expression (Bird, 2002; Schübeler, 2015). CpG island methylation has also been associated with the activity of DNA replication origins found in the vicinity of promoters (Rein et al., 1997, 1999; Martin et al., 2011). However, no single feature alone predicts the location of active origins in human cells, and GC-richness has been found to be the strongest single feature associated with active origin sites (Guilbaud et al., 2022). The reason for the preferential association of GC-rich DNA with replication origins is unclear.

In this study we investigate the epigenetic marking of active DNA replication origins, following the serendipitous discovery of short stretches of naturally dense DNA in human cells. The density of nucleic acids (measured as mass per volume) can be determined by density gradient equilibrium centrifugation in caesium salt solutions. In this physical separation technique, the opposing forces of high gravity during ultracentrifugation, and diffusion at ambient temperature, generate a stable linear gradient of caesium salts over time (Meselson et al., 1957). Any nucleic acid present in this solution equilibrates in the gradient at a density of caesium salt that corresponds to its own buoyant density. A homogeneous population of DNA forms a normal distribution in the density gradient, where the mean of the distribution corresponds to the buoyant density of the DNA, the height is proportional to the amount of DNA, and the standard deviation from the mean (0) is inversely proportional to the size of the DNA fragment, i.e. the smaller the fragment the wider the distribution (Meselson et al., 1957). Different types of nucleic acids such as RNA, RNA/DNA hybrids, single-stranded or double-stranded DNA, with their modifications, all have different characteristic densities and can be separated from each other in caesium sulphate gradients (Szybalski, 1968).

We investigated fragmented human genomic DNA by density gradient analysis and isolated and sequenced a specific DNA population with an increased mean density. This naturally dense DNA is concentrated at DNA replication origin sites in pre-replicative human quiescent and G1 phase cells, but not in replicating S phase cells. We show that the increased density of this DNA is due to an oxidation of 5-methyl-deoxycytidine. Crucially, this epigenetic marking of DNA replication origin sites before their activation is required for both DNA replication and cell proliferation.

## Results

### Isolation of naturally dense DNA from pre-replicative quiescent human cells

We synchronised human EJ30 cells in quiescence by contact inhibition and serum starvation for 12 days (Krude, 1999), purified genomic DNA from these cells, fragmented the DNA by sonication to a mean size of 250bp (range 100-400bp), and fractionated it by density equilibrium centrifugation on a caesium sulphate gradient (Figure 1A). The distribution of fragmented bulk DNA from quiescent EJ30 cells centred on a concentration of caesium sulphate with mean refractive index (RI) of 1.3683 and an overall sigma (0) of ± 0.00163. This RI corresponds to a mean density of 1.435 g/ml (Figure S1), which is consistent with the density of bulk human DNA determined previously (Guilbaud et al., 2022). However, we also observed small shoulders of lighter and denser DNA (Figure 1A). These shoulders could be due to size heterogeneity of the DNA, resulting in the superposition of narrow and wide sub-distributions with the same mean density, or they could indicate the presence of DNA fragments with different mean densities. To distinguish between these possibilities, we isolated the lighter and denser fractions and separated the DNA again on second caesium sulphate density gradients.

**Figure 1.**
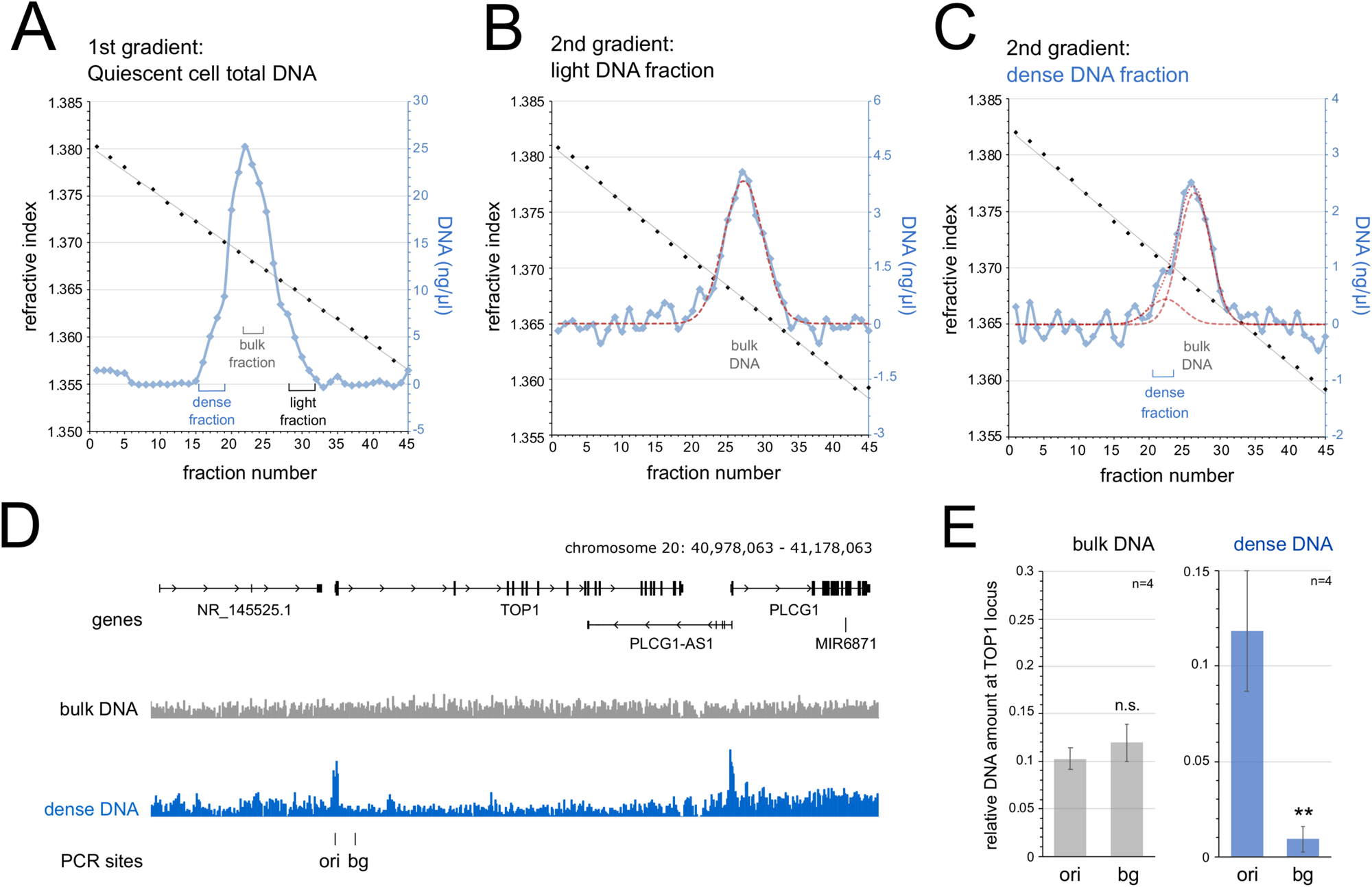
Isolation of naturally dense DNA from quiescent pre-replicative human cells. (A) Caesium sulphate density equilibrium gradient analysis of fragmented genomic DNA prepared from quiescent cell nuclei. Refractive indexes of odd-numbered gradient fractions are plotted in black together with their linear regression. DNA concentrations of each fraction across the gradient are plotted in blue after baseline and slope normalisation. Pooled DNA fractions of dense, bulk and light DNA used for loading onto second gradients and further analysis are indicated. (B) Further separation of the light DNA on a second caesium sulphate density gradient. A calculated best fit to a normal Gaussian distribution of the DNA concentrations is plotted as a hashed red line, the reference position for bulk DNA is indicated. (C) Further separation of the dense DNA on a second caesium sulphate density gradient. Calculated best fits to two normal Gaussian distributions of DNA concentrations for dense and bulk DNA are plotted by hashed (individual) and dotted (combined) red lines. Positions of pooled dense DNA fractions used further for sequencing and qPCR, and of bulk DNA are indicated. (D) Illumina sequencing read coverage profiles of isolated bulk and dense DNA at the TOP1 locus. Positions of reference genes and of sites for PCR analysis are indicated. (E) Quantitative PCR analysis of isolated bulk and dense DNA at these two sites, mean values ± standard deviations of n=4 replicates are shown (T-tests, two-tailed, unequal variance, between sites: n.s., not significant; **, p<0.01).

The fractions containing the shoulder of light DNA from the first gradient formed a distribution on the second gradient similar to bulk DNA on the first, with a mean RI of 1.3672 and a 0 of ± 0.0014 (Figure 1B). This suggests that the shoulder of light DNA on the first gradient is artefactual, and these ‘light’ fractions simply contain bulk DNA carried over from the first gradient.

In contrast, the isolated shoulder of dense DNA from the first gradient formed a biphasic distribution on the second gradient, to which two normal distributions could be fitted (Figure 1C). The major lighter distribution has characteristics similar to bulk DNA on the first gradient (mean RI = 1.3685, 0 = ± 0.0012), and it therefore corresponds to bulk DNA carried over from the first gradient. The second distribution, however, clearly showed an increased density (mean RI = 1.3706, 0 = ± 0.0013), indicating that it represents a distinct population of naturally dense DNA.

We isolated the fractions of bulk and naturally dense DNA from the first and second gradients, respectively, and used Illumina next-generation DNA sequencing to ask whether particular DNA sequences are differently represented in the two fractions. For both fractions, 126 million reads aligned uniquely to the human genome (hg38). At a representative 200 kb genomic location around the TOP1 gene locus, the aligned read coverage profiles showed strong discrete enrichment peaks for the dense DNA preparation at discrete locations, mostly in the vicinity of gene promoters present at this locus (Figure 1D). We validated this local enrichment by quantitative PCR at a peak of enriched dense DNA (TOP1 ori) and an adjacent background site (TOP1 bg). While the bulk DNA preparation showed no significant difference in abundance between these two sites, the peak site was about ten-fold overrepresented in the dense DNA compared to the background site (Figure 1E).

We conclude that we have physically isolated from quiescent human cells a distinct subpopulation of DNA fragments with an increased natural density that is strongly enriched at discrete genomic locations.

### Dense DNA is enriched at DNA replication origin sites

The enrichment site of naturally dense DNA upstream of the TOP1 gene precisely coincides with the location of a previously identified DNA replication origin (Keller et al., 2002; Langley et al., 2016), suggesting an association between dense DNA and initiation sites for DNA replication. Therefore, we compared on a genome-wide basis the locations of naturally dense DNA in quiescent cells with the locations of active DNA replication origins. We have determined these origin sites in the same human cell line previously, using the independent approaches of small nascent strand sequencing (SNSseq) (Guilbaud et al., 2022) and two variants of initiation site sequencing: IniSeq1 (Langley et al., 2016) and IniSeq2 (Guilbaud et al., 2022). To allow direct comparison with the dense DNA data, we pooled and re-aligned to the human reference genome hg38 our original raw DNA sequencing reads of two IniSeq1 datasets from EJ30 cells (Langley et al., 2016). Furthermore, we used our previously published sequencing read data from SNSseq (2-4kb length) and IniSeq2 analyses (3hr replication time), also aligned to hg38 (Guilbaud et al., 2022).

When analysed at a representative genomic locus, the discrete sites of dense DNA enrichment in quiescent cells coincide strikingly with discrete DNA replication origin sites determined by SNSseq, IniSeq1 and IniSeq2 (Figure 2). For a systematic genome-wide overlap analysis of these independent signals, we determined sequencing read enrichment peaks by the MACS2 algorithm of dense over bulk DNA. We also identified MACS2 peaks for the SNSseq, the realigned IniSeq1, and the IniSeq2 datasets to allow comparison by the same method in the same cell line. With a high stringency of peak calling (q-value of 10^-11^), we observed high genome-wide overlaps between dense DNA and DNA replication origin peaks, both visually at a representative gene locus (Figure 2A), and genome-wide (Figure 2B). These sites showed high local GC composition (Figure 2A), and about half of the dense DNA sites also overlapped with CpG islands (Figure 2B). After randomisation of the dense DNA peak locations on each chromosome, the intersects with replication origins and CpG islands were reduced by several orders of magnitude (Figure 2B), indicating that the intersects of dense DNA sites with replication origins and CpG islands are not due to chance.

**Figure 2.**
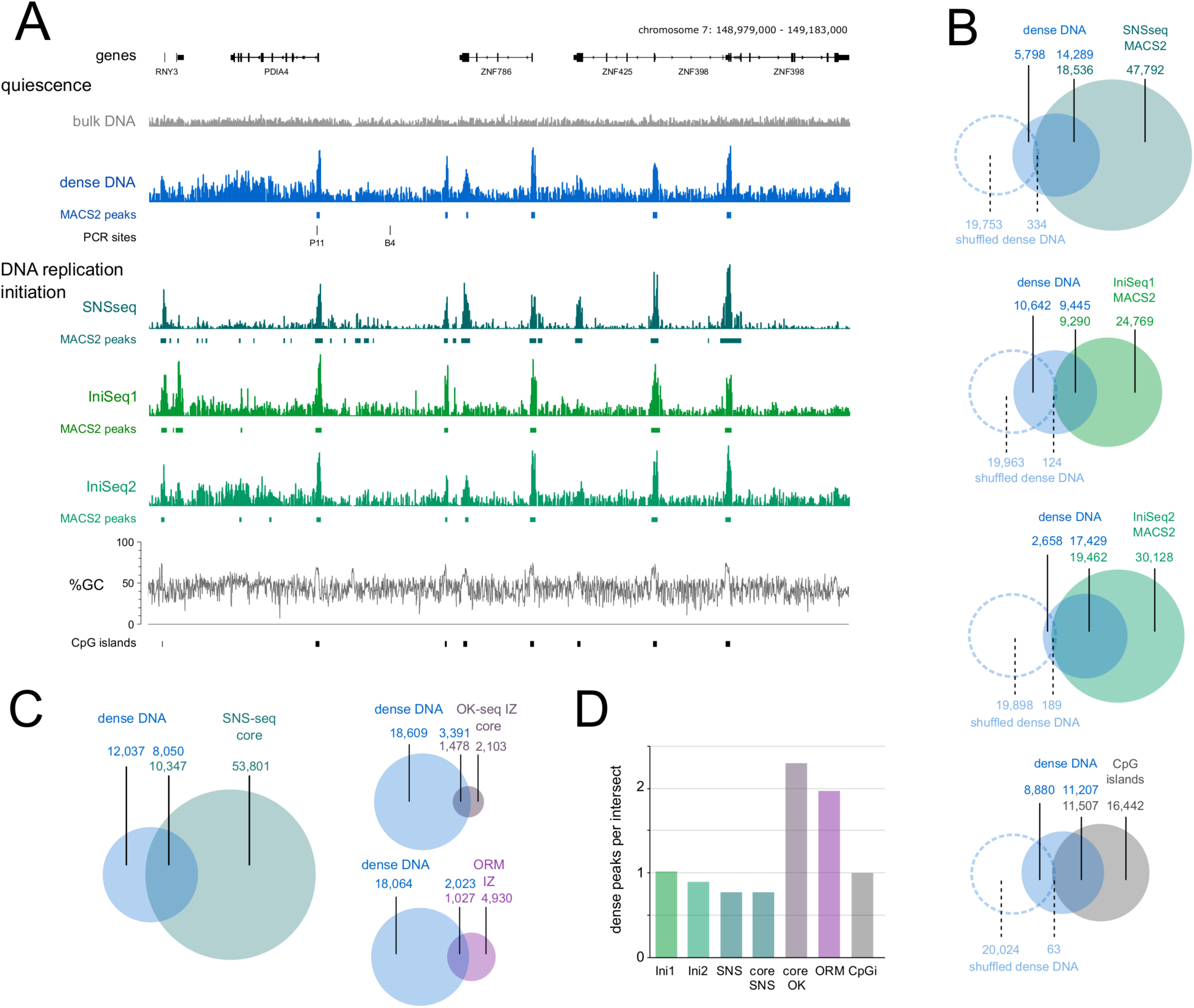
Naturally dense DNA is enriched at discrete DNA replication origins. Genome-wide analysis of dense DNA. (A) Illumina sequencing read coverage profiles at the PDIA4 locus of isolated bulk and dense DNA from quiescent cells (grey and blue), of SNSseq data (dark green) (Guilbaud et al., 2022), realigned IniSeq1 data (light green) (Langley et al., 2016), and IniSeq2 data (green) (Guilbaud et al., 2022). Positions of MACS2 peaks called with a q-value cutoff of 10^-11^ are indicated for each data set. Genome coordinates of the locus with reference genes, locations of PCR sites, percentages of GC content and CpG islands are indicated. (B) Genome-wide intersect analyses using these MACS2 peaks of dense DNA with DNA replication origin sites and CpG islands. Non-intersected and intersected numbers of peaks are indicated; the top number in the intersect indicates the number of peaks of the left distribution (dense DNA) intersecting with the right distribution (origins or CpG islands), and the bottom number indicates the number of peaks of the right intersecting with the left distribution. Intersect analyses using shuffled dense sites after randomisation per chromosome are included as hatched circles, together with corresponding peak numbers (in pale blue). (C) Genome-wide intersect analyses between dense DNA and core SNS-seq DNA replication origin sites (Akerman et al., 2020), core OK-seq initiation zones (Guilbaud et al., 2022; Petryk et al., 2016; Wu et al., 2023), and ORM initiation zones (Wang et al., 2021). (D) Average ratio of dense DNA peaks per initiation site. For each intersect, the numbers of dense DNA peaks were divided by the number of origins, initiation zones or CpG islands in the intersect.

IniSeq2 datasets allow a visualisation of origin activation by following the conversion of unreplicated template DNA to replicated, BrdU-substituted DNA over time in a cell-free system (Guilbaud et al., 2022). At representative sites of dense DNA found in quiescent cells, IniSeq2 datasets showed, after initiation of DNA replication in vitro, both a local disappearance of unreplicated template DNA and an accumulation of replicated DNA over time (Figure S2A). Dense DNA sites also overlapped substantially with IniSeq2 origins (Figure S2B) when determined by the original custom algorithm based on this conversion (Guilbaud et al., 2022). Importantly, the median peak length of replicated DNA at these dense sites increased beyond the length of the dense DNA signal over time (Figure S2C), indicating that these dense sites mark sites of active DNA replication origins, before their activation and the establishment of outwards moving replication forks.

The IniSeq2 datasets allow a ranking of origins into high, medium and low efficiency classes (Guilbaud et al., 2022). Noticeably, the high-efficiency class origins overlapped almost entirely with dense DNA sites, whilst medium and low efficiency class origins showed increasingly lower extents of overlap (Figure S2D), suggesting that dense DNA is associated preferentially with efficient origin sites.

Next, we extended the overlap analysis to different human cell lines, and to initiation zones determined by different methodologies. Dense DNA sites overlap substantially with a set of conserved SNS ‘core’ origins (Figure 2C), deduced from a panel of human cell lines (Akerman et al., 2020). OK-seq ‘core’ initiation zones deduced from OK-seq data of a panel of human cell lines (Guilbaud et al., 2022; Petryk et al., 2016; Wu et al., 2023), and initiation zones determined by ORM (Wang et al., 2021) overlapped with dense DNA sites to high and intermediate extents, respectively (Figure 2C). On average, while one dense site overlapped with one SNS-seq or IniSeq origin, two dense DNA sites were present per overlapping OK-seq or ORM initiation zone (Figure 2D).

Taken together, we conclude that naturally dense, GC-rich DNA sites in quiescent human cells show substantial genome-wide overlap with efficient DNA replication origin sites, that become activated in S phase.

### Dense DNA marks replication origins in pre-replicative cells

In our initial experiments, we identified dense DNA sequences in quiescent cells that have withdrawn from the cell cycle. We investigated next whether these dense sites persist in proliferating cells and whether they are maintained during replication origin activation *in vivo* (Figures 3 and S3). We isolated DNA from human EJ30 cells synchronised in the late G1 phase of the cell cycle by mimosine, just before initiation of DNA replication (Krude, 1999, 2000), and from cells synchronised in mid-S phase by excess thymidine, after initiation of DNA replication (Krude et al., 1997). We then separated the DNA from these cell populations on two consecutive caesium sulphate density gradients (Figure S3A, B; and Figure 3A, B). We detected dense DNA in late G1 phase cells (Figure 3A), to a similar extent (Figure 3C) and with similar characteristics (mean RI = 1.3701, 0 = ± 0.0009) to that observed in quiescent cells. In contrast, this discrete population of dense DNA was reduced to trace amounts in S phase cells (Figure 3B, C).

**Figure 3.**
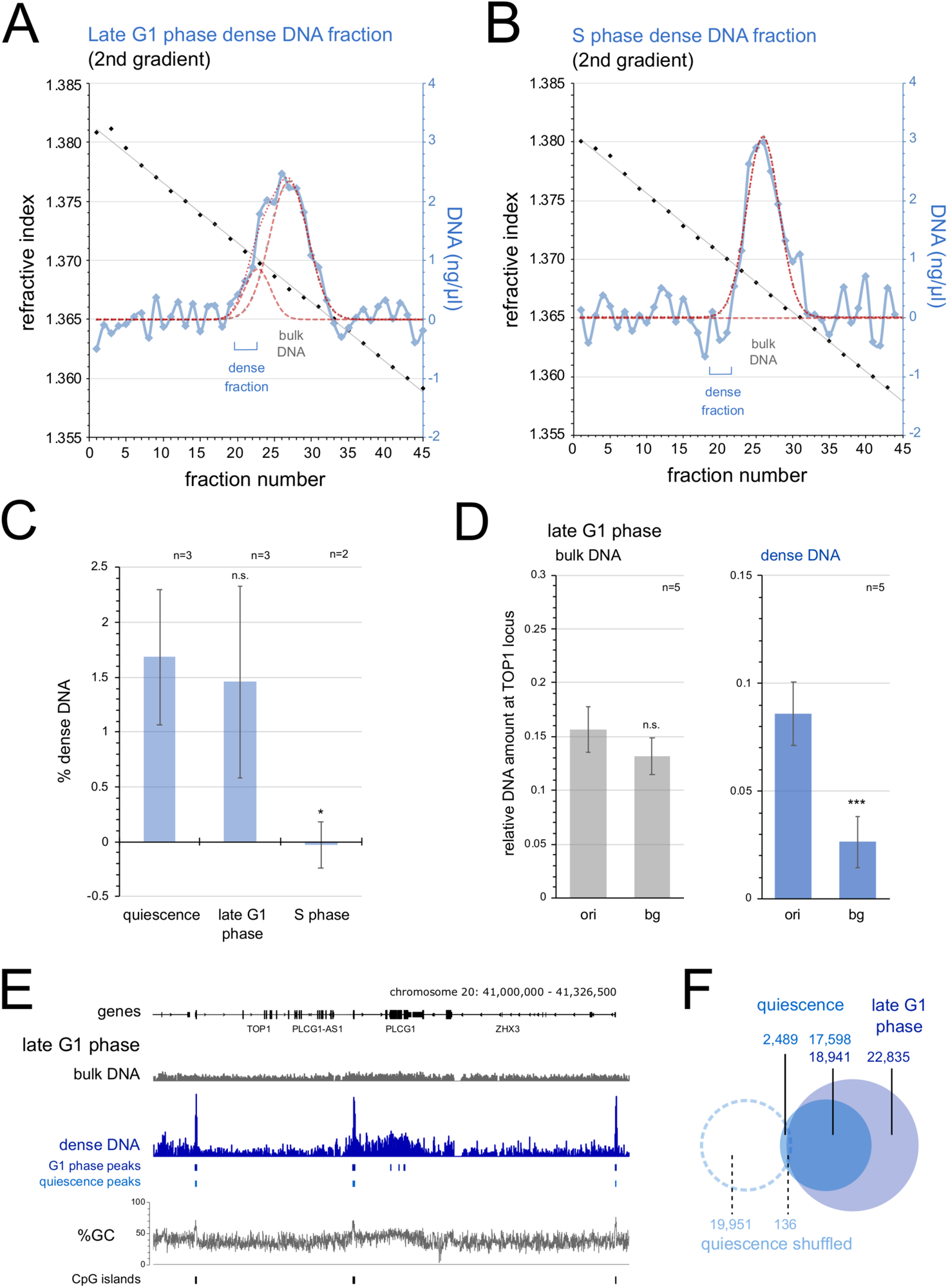
Dense origin DNA is present in late G1 phase cells and disappears in S phase. (A) Separation of dense DNA from late G1 phase cells on a second caesium sulphate density gradient. Calculated best fits of DNA concentrations to two normal Gaussian distributions are plotted by hashed (individual) and dotted (combined) red lines for dense and bulk DNA. The positions of reference bulk DNA and isolated dense DNA fractions selected for further analysis are indicated. (B) Separation of dense DNA from S phase cells on a second caesium sulphate density equilibrium gradient. Note the absence of a distinct dense DNA distribution. (C) Quantification of relative dense DNA amounts. Relative amounts were obtained as percent dense DNA of total DNA from quiescent, late G1 and S phase cells (T-tests, two-tailed, unequal variance: n.s., not significant; *, p<0.05). (D) Quantitative PCR analysis of isolated bulk and dense DNA from late G1 phase cells at TOP1 origin and background sites. Mean values and standard deviations from n=5 independent preparations are shown (T-tests, two-tailed, unequal variance: n.s., not significant; **, p<0.01; ***, p<0.001). (E) Illumina sequencing read coverage profiles at the extended TOP1 locus of isolated bulk and dense DNA from quiescent and late G1 phase cells. Positions of MACS2 peaks determined with a q-value cutoff of 10^-11^ are indicated for late G1 phase (dark blue) and quiescent (light blue) DNA samples. Genome coordinates, reference genes, a line plot for %GC content and positions of CpG islands are indicated. (F) Genome-wide intersect analysis of MACS2 peaks (q-value cutoff = 10^-11^) for dense DNA from quiescent and from late G1 phase cells (solid circles). Intersect analysis using shuffled dense sites from quiescent cells after randomisation per chromosome are included.

Quantitative PCR showed significant enrichment of the dense DNA from late G1 phase cells at the TOP1 origin over the corresponding background site (Figure 3D), as seen previously in the dense DNA from quiescent cells (Figure 1E). Next generation DNA sequencing showed enrichment of dense DNA from late G1 phase cells at the same GC-rich sites as observed in quiescent cells, seen both at a representative genomic locus (Figure 3E), and genome-wide (Figure 3F).

We conclude that the abundance of naturally dense DNA at unreplicated origin sites is found similarly in quiescent and in late G1 phase cells. In contrast, this dense DNA population is reduced to trace amounts after DNA replication origins have been activated and cells have moved into S phase.

### Dense DNA contains oxidised methyl-deoxycytidines

These observations raise the question of what DNA features cause the increase in buoyant density at DNA replication origin sites. One possibility would be the presence of non-canonic double-stranded DNA such as single-stranded DNA or RNA/DNA hybrids, which have been shown to have increased buoyant densities on caesium sulphate gradients (Szybalski, 1968). Such structures could be present in displacement loops, hairpins, G quadruplexes or R-loops, or as annealed RNA primers at replication origin sites. To address these possibilities, we degraded single-stranded nucleic acids and RNA/DNA hybrids from dense DNA preparations by single-strand specific nuclease P1 and by RNAseH, respectively. After running the treated DNA again on density gradients, we were unable to detect any significant reduction of dense DNA either at two representative origins or a corresponding background site (Figure 4A), suggesting that single-stranded domains and/or RNA/DNA hybrids do not significantly cause the increased density.

**Figure 4.**
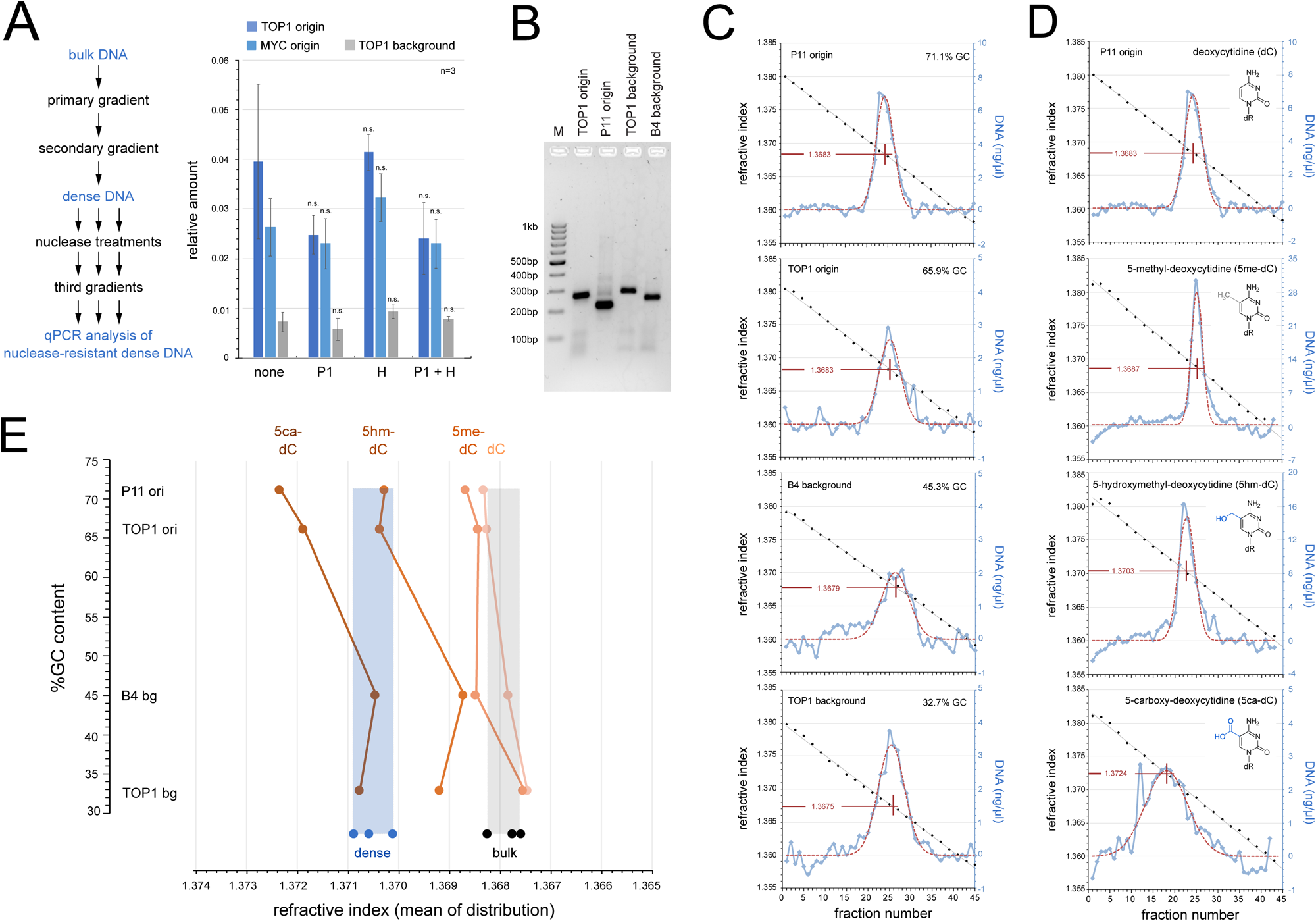
Dense DNA is a consequence of oxidised methyl-deoxycytidines. (A) Nuclease sensitivity analysis of dense DNA. An experimental flow chart is summarised on the left. Quantitative PCR analysis of nuclease-resistant, dense DNA from the third gradient is shown on the right for the TOP1 and MYC origin and the TOP background sites, mean values ± standard deviations of n=3 replicates are shown (T-tests, two-tailed, unequal variance against no nuclease input: n.s., not significant). (B) Synthesis of defined DNA fragments. PCR products of two selected origin (TOP1 and P11) and corresponding background sites (TOP1 and B4) are separated by agarose gel electrophoresis. M, 100 bp ladder. (C) Comparative density analysis of defined DNA fragments with different GC content. PCR products were synthesised using unmodified dCTP and separated on density gradients. DNA fragment identities with their %GC content are indicated for each gradient, together with fitted Gauss distributions (hatched red lines), and the refractive indexes for the means of each fitted distribution. (D) Comparative density analysis of methylated and further oxidised DNA fragments of the P11 origin site. PCR products were synthesised using unmodified and modified dCTP for an incorporation of dC, 5me-dC, 5hm-dC, and 5ca-dC. Deoxycytidine modifications are specified for each gradient. Note the increased densities for DNA fragments containing 5hm-dC and 5ca-dC, compared to unmodified or methylated dC. (E) Overview plot of mean refractive indexes against %GC content for DNA fragments with defined dC modifications. Mean refractive indexes for the indicated fragments containing either dC, 5me-dC, 5hm-dC, or 5ca-dC (shades of orange) were determined on density gradients (shown in Figs 4B, C, and S4). The ranges of mean refractive indexes of naturally dense (blue) and bulk DNA (grey) from quiescent and late G1 phase cells are superimposed as shaded boxes for reference, individual mean values determined on density gradients (shown in Figs 1, S1, 3 and S3) are plotted at the bottom of each shaded box.

The dense sites at DNA replication origins show high local GC contents that are well above the genome average of 41% and they often correspond to CpG islands (Figures 2 and 3). To exclude the possibility that GC-richness itself causes their increased density, we investigated the densities of defined double-stranded DNA fragments directly and synthesised them by PCR (Figure 4B and Table S1). We chose two GC-rich origin sites (P11 and TOP1 origin) and two associated AT-rich background sites (B4 and TOP1 background). Furthermore, we used fragment lengths for the PCR products that correspond to the average size of the fragments used for the density analyses of the natural DNA isolates (Figure 4B). The GC-rich P11 and TOP1 origin site DNA fragments equilibrated on caesium sulphate density gradients with normal distributions at mean RIs of 1.3683, and DNA fragments from the two background sites with mean RIs just below these values (Figure 4C). Therefore, the densities of unmodified DNA fragments increase very slightly and systematically with GC content (Figure 4C, E), consistent with published data (Szybalski, 1968; Panijpan, 1977). However, these slightly increased mean densities remain within the overall distribution of mean densities found for naturally bulk DNA (Figure 4E, grey box). We therefore conclude that the GC content of unmodified DNA sequence alone does not cause the formation of dense DNA at replication origin sites.

Another possibility is that covalent chemical addition of mass to the DNA would cause an increase in density. Deoxycytidine (dC) can increase in mass by chemical modification at its 5’ position (Figure 4D, inserts), first by addition of a methyl group to form 5-methyl-deoxycytidine (5me-dC) and then by oxidation to 5-hydroxymethyl-deoxycytidine (5hm-dC), 5-formyl-deoxycytidine (5f-dC), or 5-carboxy-deoxycytidine (5ca-dC) (Bird, 2002; Kriaucionis and Heintz, 2009; Tahiliani et al., 2009; Ito et al., 2011). To test this hypothesis, we synthesised by PCR the four DNA fragments with increased mass by using 5me-dCTP, 5hm-dCTP, 5f-dCTP and 5ca-dCTP and analysed their respective densities (Figure 4D, Fig S4).

Replacement of dC with 5me-dC in the P11 origin fragment resulted in a slight increase in density compared to the unmodified form (Figure 4D). A similar small increase in mean densities was seen with the other three methylated DNA fragments (Figure S4), but their values were all within the range of, or adjacent to, those of natural bulk DNA (Figure 4E, grey box). In contrast, replacement of dC with 5hm-dC caused a pronounced increase in density for all four DNA fragments (Figures 4D and S4). Importantly, this hydroxymethylation of dC shifted the densities of the two GC-rich DNA origin fragments into the range of mean densities observed for naturally dense DNA isolated from quiescent and late G1 phase cells (Figure 4E, blue box). This shift is GC-dependent as the two AT-rich background sites did not attain sufficient extra density and were positioned between the mean densities for bulk and dense DNA (Figure 4E). We were unable to synthesise any DNA containing 5f-dC, presumably because the reactive aldehyde group of 5f-dCTP would cross-link to available amines during the PCR synthesis reaction. However, replacement of dC with 5ca-dC caused a strong density shift of the two background DNA fragments into the range of naturally dense DNA and of the two origin fragments beyond that range (Figures 4D, E and S4).

We conclude that the presence of oxidised 5-methyl-deoxycytidines could explain the increased density of DNA found in pre-replicative human cells. Of all the possible oxidation states, efficient hydroxymethylation of dC would also explain the observed specificity of increased density for GC-rich origin sites, over and above the AT-rich genomic background sites.

### Naturally dense DNA at pre-replicative DNA replication origins contains 5hm-dC and 5f-dC

We investigated next whether modified deoxycytidines are present in the isolated naturally dense DNA fractions by liquid chromatography-mass spectrometry (LC-MS). We analysed nucleosides prepared from bulk and dense DNA preparations by nucleolytic degradation and dephosphorylation.

First, we calibrated the detection of mass over charge (m/z) peak values for unmodified and modified deoxycytidines after dephosphorylation of defined nucleoside triphosphates. We detected specific signals for dC, 5me-dC, 5hm-dC, 5f-dC and 5ca-dC at the expected m/z values that include H and Na as adducts (Figure S5).

The bulk DNA preparations from quiescent and late G1 phase cells contained predominantly unmodified dC, with small proportions of modified dC (Figure 5A, grey bars). In contrast, the dense DNA preparations from these cells contained predominantly 5hm-dC with substantial contributions of 5f-dC, and only small proportions of unmodified dC and 5me-dC (Figure 5A, blue bars). Significant amounts of 5ca-dC were not detected. We conclude that naturally dense DNA fragments prepared from quiescent and late G1 phase human cells contain predominantly oxidised forms of 5-methylcytidine, namely 5hm-dC and, to a lesser extent, 5f-dC. Next, we asked if 5hm-dC occurs at DNA replication origin sites in dense DNA preparations. The restriction endonuclease MspI cuts DNA at CCGG sites regardless of whether the dC residues at this site are unmodified, methylated or hydroxymethylated (Figure 5B, top). However, glucosylation of 5hm-dC residues at their 5-hydroxyl group by T4 beta-glucosyltransferase (T4-BGT) renders the CCGG site resistant to cleavage by MspI (Figure 5B, bottom). The DNA sequence of the GC-rich TOP1 DNA replication origin fragment contains two adjacent MspI sites (Table S1), allowing an analysis of this site by MspI resistance followed by PCR (Figure 5B). We therefore glucosylated bulk and dense DNA preparations by T4-BGT, digested the DNA with MspI, and determined by PCR whether the TOP1 origin site had been cut, and therefore contained 5hm-dC (Figure 5C-E). A correctly sized 265bp TOP1 origin PCR product was synthesised from uncut bulk and dense DNA fractions of quiescent cells (Figure 5C), as expected. Prior digestion of these DNA fractions with Msp1 prevented synthesis of the PCR product. Importantly, treatment with T4-BGT rendered a substantial fraction of the dense, but not the bulk DNA fraction resistant to digestion by MspI yielding a defined PCR product, indicating that the dense TOP1 origin site contains 5hm-dC (Figure 5C, bottom panel).

**Figure 5.**
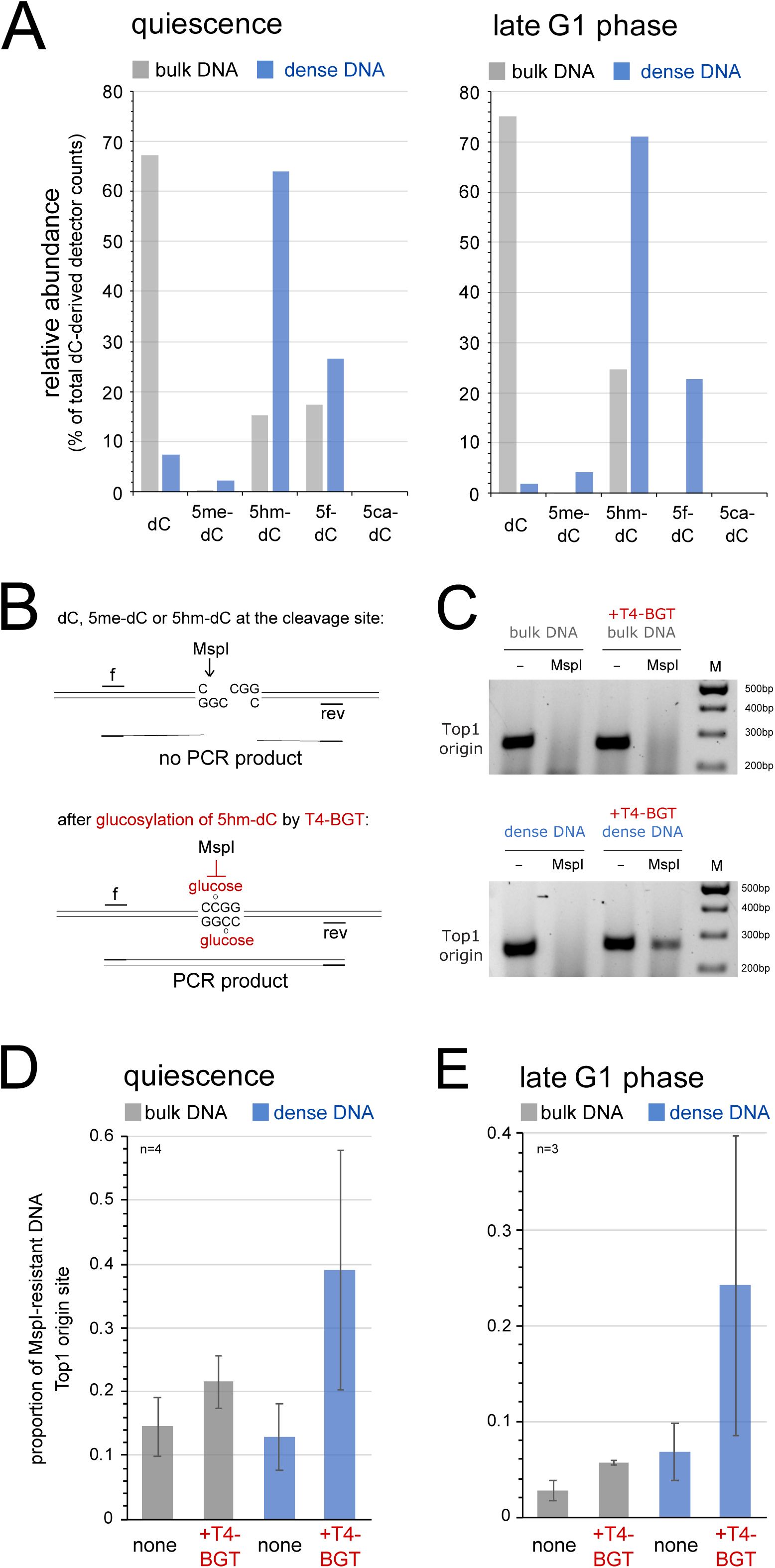
Dense origin DNA contains predominantly 5hm-dC and 5f-dC. (A) Mass spectrometry analysis of relative proportions of deoxycytidine-derived nucleosides in the bulk and dense DNA fractions from quiescent and late G1 phase cells. LC-MS detector counts for the individual dC-derived nucleosides are plotted as the percentages of the total counts from all dC-derived nucleosides. (B) Principle of specific detection of 5hm-dC by a glucosylation-dependent sensitivity assay to the restriction endonuclease MspI. Glucosylation of 5hm-dC via the 5-OH group by T4 beta-glucosyltransferase (T4-BGT) renders the restriction site CCGG resistant to MspI, while native dC, 5me-dC and 5-hm-dC are cleaved. Msp-resistant DNA at target origin sites is detected by PCR. (C) PCR product analysis. Bulk and dense DNA fractions from quiescent EJ30 cells were glucosylated by T4-BGT and subsequently digested with Msp1 as indicated. DNA samples were amplified by PCR using the TOP1 origin primer pairs and products analysed by agarose gel electrophoresis. (D-E) Quantitative PCR analysis. Bulk and dense DNA samples from quiescent and late G1 phase cells were amplified by quantitative PCR at the TOP1 origin site after glucosylation by T4-BGT and MspI cleavage. Results are expressed as proportions of Msp1-resistant DNA compared to the corresponding uncut samples. Mean values and standard deviations of n=3-4 technical replicates are shown.

We then used quantitative PCR to determine the extent of 5hm-dC at the TOP1 origin site in DNA preparations from quiescent and late G1 phase cells using the MspI sensitivity assay (Figure 5D, E). Glucosylation of 5hm-dC by T4-BGT caused only a marginal increase of MspI-resistant DNA over background levels in the bulk DNA fractions, whereas a four-fold increase was observed in the dense DNA. We conclude that dense DNA contains 5hm-dC at both CCGG sites at the TOP1 DNA replication origin in quiescent and late G1 phase cells.

Taken together, these data show that DNA origin sites predominantly contain the oxidised methyl-deoxycytidines 5hm-dC and 5f-dC in pre-replicative quiescent and late G1 phase cells before DNA replication initiates at these sites in S phase. These modified deoxycytidines cause an increase in the natural density of the DNA at these origin sites, allowing their biophysical isolation from bulk DNA.

### Dense DNA at replication origins depends on DNMT1 and TET enzyme activities

Finally, we investigated whether biosynthesis of oxidised methyl-deoxycytidines at DNA replication origins is required for DNA replication and cell proliferation.

The biogenesis of oxidised 5me-dC in human cells occurs in two enzymatic steps following DNA replication and deposition of unmodified dC by DNA polymerases. Each step can be blocked by specific inhibitors (Figure 6A). First, DNA methyltransferases (DNMT) catalyse the methylation of dC into 5me-dC (Bird, 2002). Secondly, Ten-Eleven-Translocation DNA dioxygenases (TET1-3) catalyse the oxidation of 5me-dC into 5hm-dC, and then on to 5f-dC and 5ca-dC (Tahiliani et al., 2009; Ito et al., 2011; Zhang et al., 2023). Specific small molecule inhibitors are available to block the activities of both DNMT and TET enzymes (Figure 6A), and we used these to evaluate any functional role of the biogenesis of oxidised forms of 5me-dC for DNA replication and cell proliferation.

**Figure 6.**
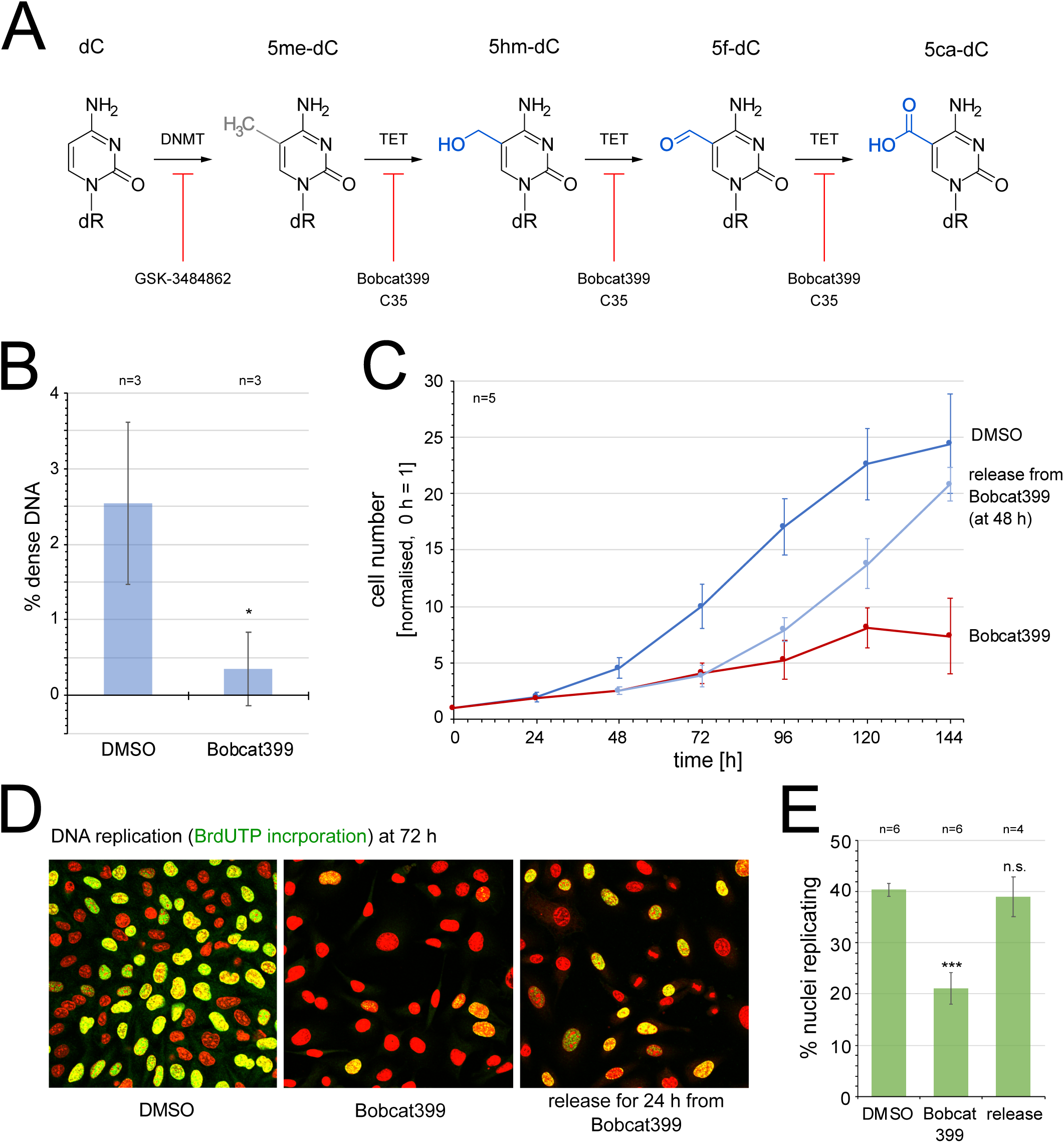
Oxidation of methyl-deoxycytidine at DNA replication origins is required for cell proliferation and DNA replication. (A) Overview of 5-methylation and further oxidation of deoxycytidine by DNA methyltransferase (DNMT) and Ten-eleven translocation DNA dioxygenase (TET) enzyme activities. Specific inhibitors of DNMT and TET enzyme activities are indicated. (B) Quantification of relative amounts of dense DNA in proliferating EJ30 cells treated for 72 hours with either DMSO or 125 µM of the TET inhibitor Bobcat399. Relative amounts were obtained as percentage dense DNA of total DNA. Mean values and standard deviations from n=3 independent gradient analyses are shown (T-test, two-tailed, unequal variance: *, p<0.05). (C) Inhibition of cell proliferation. Asynchronously proliferating EJ30 cells were treated with DMSO (blue) or 125 µM Bobcat399 (red) for 144 h. For release, Bobcat399 was replaced by DMSO in the culture medium after 48 h and cultivation continued for further 96 h (pale blue). Normalised cell numbers are plotted against time as averages ± standard errors of the mean for n=5 independent experiments. (D, E) Inhibition of DNA replication. After 72 h of treatments, replicating cells were pulse-labelled with BrdU and replicating cell nuclei detected by confocal immunofluorescent microscopy. (D) Representative micrographs. DNA is stained with propidium iodide (red), BrdU incorporation with specific antibodies (green). Note the lower proportions of replicating cell nuclei and lower intensity of BrdU incorporation signals in Bobcat399-treated cells compared to the DMSO control and the release. (E) Quantification of the percentages of replicating cell nuclei. Proportions of nuclei replicating their DNA were scored and plotted for the three growth conditions. Mean values ± standard errors of the mean from n=4-6 independent preparations are shown (T-tests, two-tailed, unequal variance between DMSO control and treatments: n.s., not significant; ***, p<0.001).

Bobcat399 is a cytosine-based compound that inhibits TET enzyme activity competitively and leads to a significant reduction of intracellular 5hm-dC (Chua et al., 2019). Addition of Bobcat399 to proliferating human cells resulted in a ten-fold reduction in naturally dense DNA in these cells after 72h of treatment, as determined by density gradient centrifugation (Figures 6B and S6A, B). Therefore, the synthesis of naturally dense DNA in human cells in vivo depends on the oxidation of 5me-dC residues by TET enzyme activity.

We next analysed the requirement of TET enzyme activity for cell proliferation and DNA replication. Addition of Bobcat399 to proliferating human cells resulted in a marked reduction in cell proliferation at 48 hours of treatment and led to cytostatic arrest after several days (Figure 6C). Concomitant with an inhibition of cell proliferation, inhibition of TET enzyme activity also resulted in an inhibition of DNA replication. After 72 hours of treatment with Bobcat399, about half as many cells incorporated BrdU as untreated control cells, and the extent of BrdU incorporation was much lower in the TET enzyme-inhibited cell nuclei (Figure 6D, E). Importantly, this inhibition is reversible as cells resumed efficient DNA replication within 24 hours, and proliferation within 48 hours after removal of Bobcat399 (Figure 6C-E), indicating that DNA replication and cell cycle progression can resume once the epigenetic marking of DNA by 5me-dC oxidisation is regenerated.

Next, we validated these findings by using an independent small molecule inhibitor of TET enzyme activity. The cell-permeable compound 35 (C35) targets the catalytic core of TET enzymes and inhibits their enzymatic activities allosterically (Singh et al., 2020). As with Bobcat399, we found that treatment of proliferating human cells with C35 resulted in a cytostatic arrest of cell proliferation and an inhibition of DNA replication (Figure S6C-E). This sensitivity to two unrelated inhibitors suggests strongly that the biogenesis of 5hm-dC via TET-dependent oxidation of 5me-dC is required for cell proliferation and DNA replication.

A prediction of this finding is that inhibiting the prior methylation of dC to 5me-dC should also have a negative effect on cell proliferation and DNA replication, because 5me-dC is the substrate for TET enzyme activity. GSK-3484862 is a non-covalent small molecule inhibitor of DNMT1 activity, which also leads to degradation of DNMT1 and to subsequent DNA hypomethylation in human cells (Pappalardi et al., 2021; Chen et al., 2023). Consistent with this prediction, we found that addition of GSK-3484862 to proliferating human cells resulted in: (i) a reduction of naturally dense DNA, (ii) a cytostatic inhibition of cell proliferation and (iii) a significant inhibition of DNA replication (Figure S6F-L).

Taken together, our data show that the biogenesis of oxidised forms of 5me-dC via DNMT and TET enzyme activities results in an accumulation of naturally dense DNA at human DNA replication origin sites before their activation. Importantly, this biogenesis of oxidised forms of 5me-dC is functionally required for DNA replication and cell proliferation. We conclude that sites of initiation of DNA replication are defined epigenetically by oxidation of 5me-dC to 5hm-dC and 5f-dC.

## Discussion

Here, we have isolated from human cells a distinct DNA population with a higher natural density than bulk DNA. Genome-wide analyses established that this dense DNA is specifically enriched at discrete sites of the genome that correspond to DNA replication initiation sites identified in the same cell line by SNS-Seq (Guilbaud et al., 2022) and Ini-Seq1 (Langley et al., 2016) and in particular, to efficient origins identified by Ini-Seq2 (Guilbaud et al., 2022). They also overlapped with conserved core initiation sites and initiation zones identified in different cell lines by additional approaches. Using cell cycle analyses, we established that this dense DNA is present in quiescent and in late G1 phase cells prior to DNA replication, but absent from S phase cells after the onset of DNA replication. These observations indicate that dense DNA is converted to less dense DNA by DNA replication and that the density is restored again by the time cells reach late G1 phase in the subsequent cell cycle, or they enter quiescence. We then identified the underlying DNA modifications that render DNA naturally dense as oxidised forms of 5-methyl-deoxycytidine (5hm-dC and 5f-dC), using four separate lines of investigation: (i) calibration of density gradients with DNA fragments of defined GC content and defined dC modifications, (ii) mass spectrometry, (iii) modification specificity of the restriction endonuclease MspI, and (iv) inhibition of biosynthetic pathways of dC methylation and further 5me-dC oxidation. Importantly, these latter inhibition studies established that the generation of oxidised methylcytidine at DNA replication initiation sites is required for DNA replication and cell proliferation.

### Oxidised methyl-dC at DNA replication origin sites

The oxidative modification of 5-methylated cytosine residues in DNA to 5-hydroxymethyl, 5-formyl-, and 5-carboxyl-dC by TET enzymes was first described some fifteen years ago (Kriaucionis and Heintz, 2009; Tahiliani et al., 2009; Ito et al., 2011). With the development of high resolution and specific detection methods, 5hm-dC and 5f-dC were found to be stable components of DNA, as opposed to transient metabolic products, and they were present in all tissues and cell types investigated (Globisch et al., 2010). We show here that the presence of oxidised 5-methyldeoxycytidines increases the buoyant density of GC-rich DNA, presumably by the increase in molecular mass associated with the additional oxygen moiety. We exploited this property to isolate and further analyse this dense DNA, and show that oxidised forms of 5-methyl-deoxycytidine, namely 5hm-dC and 5f-dC, coincide with discrete start sites of DNA replication, before DNA replication is initiated from these sites.

An association of methylated DNA with replication origin sites has been suggested before. In the pre-genomic era, a high concentration of methylated CpG dinucleotides were found by bisufite sequencing at several candidate vertebrate DNA replication origin loci known at the time, and methylation correlated with origin activity at these sites (Rein et al., 1997, 1999). In an early genome-wide SNS-seq analysis, active replication origins were found associated with methylated CpG islands, and the extents of nascent strand enrichment and CpG methylation correlated with each other (Martin et al., 2011). Importantly, methylation data derived from bisulfite sequencing methodology used at that time do not differentiate between the presence of 5me-dC or 5hm-dC (Huang et al., 2010). Therefore, this reported association of ‘methylated’ CpG islands with active DNA replication origins (Martin et al., 2011) is not inconsistent with the marking of active origins by oxidised 5me-dC, as we report here. The presence of 5me-dC, 5hm-dC and 5f-dC at CpG islands has been shown subsequently in mouse embryonic stem cells (mESCs) by the development of bisulfite sequencing methodologies that differentiate between these variants (Song et al., 2011; Booth et al., 2012; Song et al., 2013; Shen et al., 2013; Booth et al., 2014). Although no association with replication origins was investigated in these studies, the high extent of overlap between replication origins and modified CpG islands is consistent with an accumulation of these modified dC residues at replication origins. In a recent report (Prikrylova et al., 2019), early SNS-ChIP data from mESCs (Cayrou et al., 2011) were intersected bioinformatically with early DNA immunoprecipitation and sequencing (DIP-seq) datasets, obtained with antibodies specific to 5me-dC and 5hm-dC (Shen et al., 2013). The SNS data intersected four times more with 5hm-dC than with 5me-dC DIP-seq sites, suggesting a preferred association of replication origins with 5hm-dC (Prikrylova et al., 2019). Therefore, the marking of active human DNA replication origins with oxidised 5me-dC reported here is supported by earlier studies in vertebrate cells.

### Cell cycle dynamics and biogenesis of oxidised methyl-dC

We detected dense DNA in pre-replicative quiescent and late G1 cells, but not in replicating S phase cells (Figures 1-3). This observation is consistent with genome-wide data on the re-establishment of dC methylation and hydroxymethylation after DNA replication. DNA methylation is predominantly symmetric on both DNA strands at CpG sites and newly replicated hemi-methylated dC becomes fully methylated again in replication-coupled and independent pathways (Petryk et al., 2021). Re-methylation of dC is not immediate and the time taken for efficient restoration ranges from a few minutes up to several hours in different human and rodent cell types, and these times also vary with the experimental methods employed (Rein et al., 1999; Charlton et al., 2018; Xu and Corces, 2018; Ming et al., 2020; Stewart-Morgan et al., 2023). Most of the replicated DNA is methylated again when a cell reaches mitosis, but complete restoration of genome methylation may take place afterwards during the subsequent cell cycle. In contrast, hydroxymethylation of dC is slower. A recent work using a quantitative mass spectrometry approach for detecting DNA modifications on maturing nascent DNA in mouse embryonic stem cells (iDEMS) concluded that dC hydroxymethylation kinetics of nascent DNA lag behind those of dC methylation genome-wide, and take several more hours to reach pre-replicative levels again (Stewart-Morgan et al., 2023). Genome-wide, 5hm-dC is a very stable mark, and in proliferating cells it is predominantly associated with parental rather than nascent DNA strands (Bachman et al., 2014; Prikrylova et al., 2019; Stewart-Morgan et al., 2023). Taken together, these data suggest that 5hm-dC is stably inherited on the parental DNA strands and that newly synthesised, initially unmodified, DNA strands become methylated and then oxidised following DNA replication.

Inhibition of DNMT and TET enzyme activities by three unrelated small molecule inhibitors all resulted in an inhibition of both DNA replication and cell proliferation (Figures 6, S6). These observations imply a functional requirement for the synthesis of oxidised 5me-dC for origin activation and the initiation of DNA replication. However, overexpression experiments using active or even inactive catalytic domains of TET2 in HeLa cells resulted in a cell cycle delay, leading the authors to conclude that 5hm-dC acts as a barrier to DNA replication (Prikrylova et al., 2019). Our data reported here do not support this conclusion. It may be that the cell cycle delays observed by Prikrylova and colleagues are due to dominant-negative effects caused by the overexpressed active or inactive TET2 protein domains.

The inhibition of cell proliferation by DNMT and TET inhibitors reported here is of potential clinical importance. It suggests that existing anti-cancer treatment strategies targeting DNA methylation (Lee et al., 2024), which are thought to inhibit cell proliferation indirectly via transcriptional regulation of oncogenes and tumour suppressors, may also execute anti-proliferative effects more directly by interfering with DNA replication. Further work in this direction should therefore be encouraged.

### An epigenetic model for the specification of human DNA replication origins

Taking these findings together, we suggest a possible mechanism for how human DNA replication origins are marked epigenetically by dC methylation and subsequent oxidation (Figure 7). First, at the level of primary genetic encoding, human DNA replication origins are generally rich in GC base pairs and often, but not necessarily always, coincide with CpG islands or contain G quadruplexes (Martin et al., 2011; Besnard et al., 2012; Valton et al., 2014; Picard et al., 2014; Langley et al., 2016; Prorok et al., 2019; Akerman et al., 2020; Guilbaud et al., 2022; Hu and Stillman, 2023). Second, an initial epigenetic mark of dC methylation is added to these GC-rich origin sites by DNMT activity. This first-level modification of GC-rich origin DNA only leads to a slight increase in density and it serves as an intermediate. Third, methylated GC-rich origin sites become substrates for subsequent oxidation by TET enzyme activity. These oxidised methylcytidines now significantly increase the buoyant density of the GC-rich DNA origin fragments, which has allowed us to isolate them from pre-replicative human cells by density equilibrium centrifugation. Crucially, they are also required for the initiation of DNA replication.

**Figure 7.**
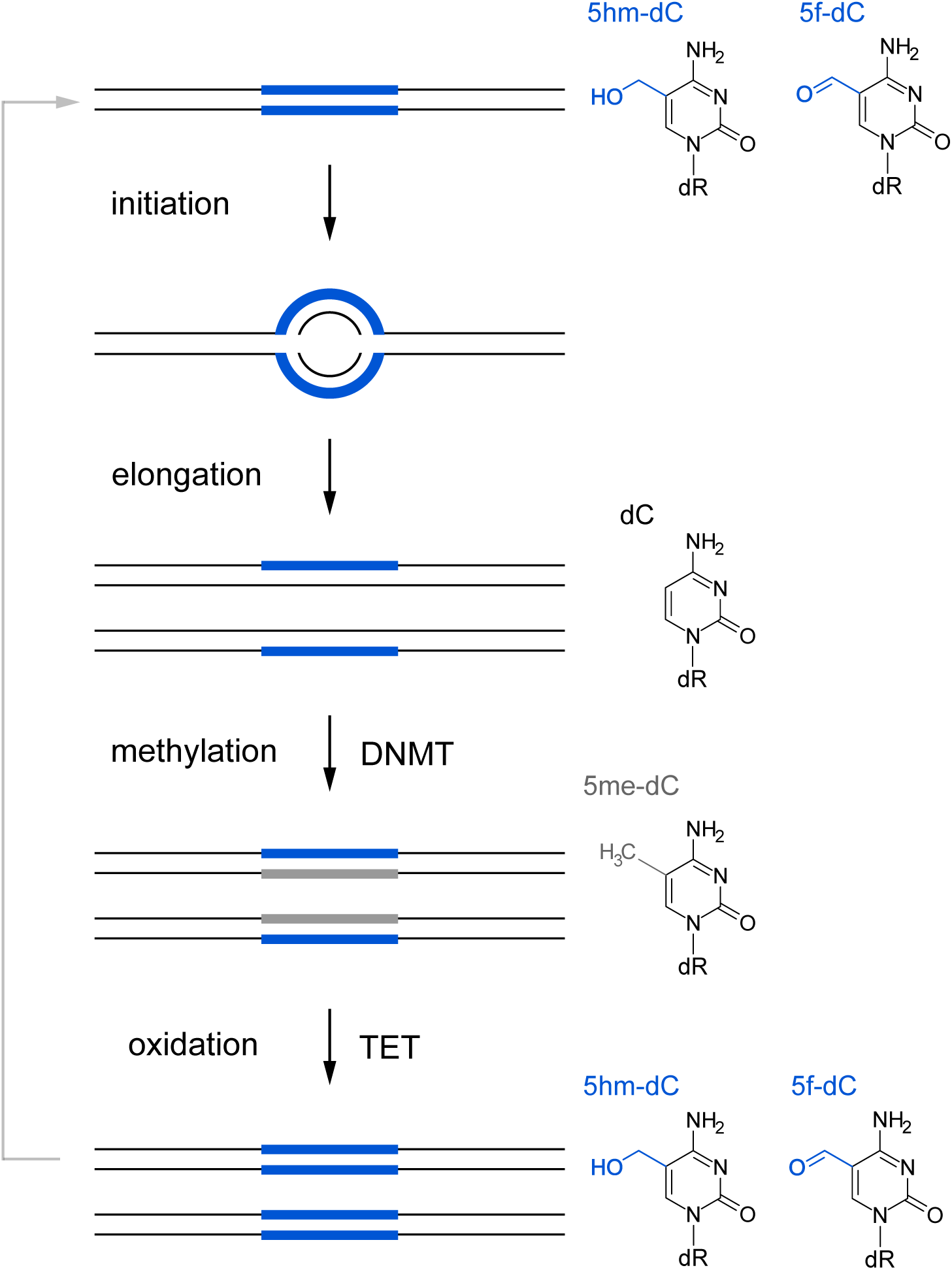
Model depicting a mechanism for the epigenetic specification of DNA replication origins by deoxycytidine methylation and further oxidation during a DNA replication cycle. Bulk DNA strands are indicated by thin black lines, and epigenetically marked DNA sites are indicated by wide lines. Cytidine modifications conferring substantially increased density are indicated in blue, modifications not substantially increasing density are indicated in grey. The underlying chemical modifications are specified on the right, using the same colour-coding. DNMT, DNA methyltransferases; TET, Ten-eleven translocation DNA dioxygenases.

This genetic and epigenetic marking of origin sites can be integrated with the semiconservative DNA replication cycle (Figure 7). Newly replicated, nascent DNA strands emanating from an activated origin contain unmodified bulk cytidines. The parental strands containing 5hm-dC (and 5f-dC) maintain their epigenetic marks and are paired with unmodified nascent strands. This semiconservative inheritance of marked parental DNA strands results in a significant reduction in associated density at the replication initiation site, consistent with a lack of dense DNA in S phase (Figure 3) and the lower density associated with reduced modified GC content (Figure 4). Subsequently, the unmodified nascent strands become substrates for DNMT-dependent methylation (Figure 7). However, methylation does not increase the density of origin DNA substantially and this hybrid site would therefore still have reduced density compared to the high-density levels observed before replication (Figure 4). High GC content and dC methylation could therefore be considered necessary, but not sufficient, to mark DNA replication origins. Full acquisition of high density and epigenetic marking of the DNA site would finally be achieved by oxidation of methylated dC by TET enzyme activity, expected to be completed after DNA replication. Once oxidation is complete, the cycle is reset and the DNA replication origin is rendered dense again and marked for the subsequent replication cycle (Figure 7).

This semiconservative model can therefore explain how a defined origin site marked for initiation of DNA replication is maintained during the cell cycle from one generation to the next (Figure 7). It also provides a new mechanistic explanation for DNA replication licensing to restrict initiation of DNA replication at each specified origin to once per cell cycle (Laskey et al., 1981; Blow et al., 1987). In this interpretation, a license would be provided in cis by oxidised methylcytidines on both DNA strands to specify origins before their activation. This specification is removed on the nascent strands by the process of replication and is not restored until S phase is complete and the cell has entered the next cell cycle.

Finally, this epigenetic model (Figure 7) also allows for flexibility in origin site specification and for dynamic reprogramming. For an existing initiation site to be silenced, the accessibility of newly replicated chromatin to DNMT or TET enzymes could be restricted, perhaps by chromatin remodelling, so that nascent strands do not receive the mark while the parental mark is inherited and eventually diluted out during subsequent cell cycles. Alternatively, oxidised methylcytidines could be removed by base excision repair involving thymine DNA glycosylase (TDG) (He et al., 2011; Wu and Zhang, 2017). Either would return the site to an epigenetically unmarked, inactive, but yet genetically defined GC-rich default state. In turn, an unmarked GC-rich site could be selected as a new DNA replication initiation site by recruitment of DNMT and TET activities to generate a high density of oxidised methylcytidines de novo.

Currently, we do not know how this epigenetic specification of DNA replication origin sites leads to the initiation of DNA replication at this site. It is reasonable to assume that these modifications may increase the probability for recruiting replication initiation factors to origin sites, in analogy to the binding of the origin recognition complex ORC to DNA sequence-specified origins in budding yeast (Hu and Stillman, 2023). Future work will be required to address this exciting perspective.

## Materials and Methods

### Cell culture

Human male EJ30 bladder carcinoma cells (Hastings and Franks, 1981) were cultured as proliferating monolayers as described (Krude, 1999).

Cells were synchronised in quiescence by contact inhibition and serum starvation in late G1 phase by a 24 h treatment with mimosine (0.7 mM; Sigma), and in S phase by a 24 h treatment with excess thymidine (2.5 mM; Sigma), as described (Krude, 1999; Krude et al., 1997). Synchronisation was confirmed by flow cytometry of isolated nuclei and staining of DNA content with propidium iodide as described (Krude et al., 1997), using a CytoFLEX cytometer (Beckman Coulter).

For proliferation assays, cells were seeded at low density onto 13 mm glass coverslips and propagated in 24-well dishes (Nunc) containing 1 ml growth medium. For inhibition of DNMT and TET enzyme activities, the following inhibitors were dissolved in DMSO: GSK-3484862 (ChemieTek, at 10 mM), Bobcat339 (Sigma, at 16.8 mM), C35 (AOBIOUS, at 20 mM), and added to growth media at final concentrations of 10 µM, 125 µM and 150 µM, respectively. Controls contained the corresponding volumes of DMSO only. For monitoring release from inhibitor treatment, growth medium was replaced at 48 h with medium from untreated cells containing DMSO only.

For quantifying relative cell numbers, coverslips were taken at 24 h intervals, washed in PBS, fixed for 5 min in para-formaldehyde (4%), washed in PBS, stained for 5 min with DAPI (Sigma) at 1 μg/ml in PBD and 0.5% Triton X-100, and mounted onto microscope glass slides in 70% glycerol / 30% PBS. Coverslips were imaged using a Zeiss Axioskop 40 fluorescence microscope (20x objective lens) fitted with a Retiga R1 greyscale camera (QImaging). Ten images were taken of sequential, non-overlapping fields of view per coverslip and the numbers of cells per image were counted. Mean cell counts were normalised to mean cell counts at the start of each experiment.

For DNA replication assays, cells growing on glass coverslips were labelled with 5-bromo-2’-deoxyuridine (BrdU, 33 μM) for 45 min before fixing in ice cold methanol and storage at -20°C. For staining, coverslips were rehydrated stepwise in decreasing concentrations of methanol (from 100% to 0% in 20% intervals, 10 min each at room temperature), treated with 1 M HCl for 1 h, washed in H_2_O and PBS, and incubated in blocking buffer (3% dried milk powder, 0.1% Triton X100, 0.02% SDS in PBS) for 30 min. Immunostaining was performed for 1 h using primary mouse anti-5-bromodeoxyuridine antibody (Sigma) at 2-5 µg/ml, secondary Alexa488-conjugated goat anti-mouse antibody (Molecular Probes) at 2 µg/ml, and propidium iodide (Sigma) at 20 µg/ml, all in blocking buffer. Imaging was carried out with an Olympus FLUOVIEW FV3000 confocal microscope (60x objective), using 561 nm and 488 nm lasers for propidium iodide (red channel) and BrdU (green channel,) respectively. Images were processed in FIJI (ImageJ) and cells were scored manually as replicating (red and green) or non-replicating (red).

### Preparation of genomic DNA

Cell nuclei were isolated by hypotonic treatment, Dounce homogenisation and centrifugation as described (Krude, 1999, 2000). The pelleted nuclei were resuspended and washed in DNA buffer (10 mM Tris-Cl, pH8.0; 125 mM NaCl; 1 mM EDTA), pelleted again, dissolved in 750 µl lysis buffer (10 mM Tris-Cl, pH8.0; 125 mM NaCl; 1 mM EDTA, 1% l-laurylsarkosine, 2mg/ml proteinase K) and incubated at 55°C for 24 h. High-molecular weight DNA was extracted with phenol/chloroform, precipitated with ethanol and dissolved in DNA buffer at 55°C for 3 h. Concentrations were determined by spectrophotometry using a NanoDrop 1000 spectrophotometer (Thermo Fisher Scientific).

For DNA fragmentation, 20 µg of high molecular weight DNA per sample were adjusted to a 130 µl volume in DNA buffer and fragmented on a Covaris focussed ultrasonicator ME220 (using microTUBE-130 AFA Fiber Strips V2 and the following settings: 130 s duration, 70 W peak power, 20% duty factor, 1,000 cycles per burst, average power of 14.0 at 20°C). Two samples were processed in parallel for each reaction, and subsequently pooled. Resulting distributions of fragmented double-stranded DNA were checked to a target size of 250 bp by neutral agarose gel electrophoresis.

### Equilibrium density gradient centrifugation

DNA samples were adjusted to 8 ml of 1.5 M Cs_2_SO_4_, 10 mM Tris-Cl pH 7.4, 1 mM EDTA (corresponding to a refractive index of 1.3700 at 20°C) and loaded into polypropylene bell-top Quick-Seal tubes (Beckman Coulter). Centrifugation was performed in a near-vertical MLN-80 rotor (Beckman Coulter) at 60,000 rpm at 20°C for 20 h in an OptimaMAX-XP ultracentrifuge (Beckman Coulter). Gradients were pumped out from bottom to top through a glass capillary tube attached to silicone tubing, using peristaltic pump P-1 (GE Healthcare) at a flow rate of 2 ml/min. Fractionation was performed manually at 8 drops per fraction at 20°C.

Refractive indexes of odd-numbered fractions were determined with an ATAGO R5000 hand refractometer. Densities were determined gravimetrically by weighing a 100 µl volume in a sealed container at 20°C on Sartorius SECURA26-1S precision scales. Scales and pipettes were calibrated to the density of distilled water (1 g/ml) at 20°C. To convert refractive indexes (RI) to densities (D), a linear best fit was determined experimentally (D = 12.46g/ml x RI -15.62g/ml, N = 38, R^2^ = 0.9945; Fig. S1).

DNA concentrations of every fraction were determined by spectrophotometry using a NanoDrop 1000 spectrophotometer (Thermo Fisher Scientific). Samples were blanked against the caesium sulphate solution used for the gradients (1.5 M Cs_2_SO_4_, 10 mM Tris-Cl pH 7.4, 1 mM EDTA). Averages of at least four readings were taken for each fraction.

Raw DNA measurements showed a slight linear baseline tilt from high to low densities. Measurements were therefore corrected for slope and intersect of the baseline readings for each individual gradient. A linear best fit regression was calculated from the measurements of fractions not containing RNA or DNA (i.e. excluding fractions with RI > 1.3780; and RI between 1.365 and 1.372). For each fraction, the slope and intersect values derived from the best-fit line were subtracted from the actual measurement values. This correction resulted in normalised values of DNA concentrations for all fractions, with baseline slope and intersect values of zero. Fits to Gaussian distributions were calculated for DNA concentration profiles, after baseline correction, by minimising the sums of squared differences between the experimental data and the reference Gaussian distributions.

For each experiment two consecutive gradients were run to increase removal of contaminating light bulk DNA from the denser DNA fractions. Raw DNA preparations were separated on a primary density gradient. Fractions of primary gradients containing bulk DNA (RI = 1.3670 – 1.3682) were isolated and pooled. Fractions of the primary gradients covering the area of dense DNA (RI = 1.3695 – 1.3715), and of the area covering light DNA (RI = 1.3630 – 1.3660), were collected and run again on secondary density gradients. Fractions of the secondary gradients containing dense DNA were isolated and pooled.

Proportions of dense over total DNA were calculated after baseline correction by dividing the integrated DNA amounts of an extended dense area of the secondary gradient (between RIs of 1.3700 and 1.3750) by the integrated DNA amounts of the total DNA from the corresponding primary gradient.

### Further analysis of dense and bulk DNA fractions

DNA from the isolated and pooled fractions was desalted on PD MiniTrap G-25 columns (GE Healthcare) equilibrated in 10 mM Tris-Cl pH 7.4, 1 mM EDTA.

For a degradation of single-stranded DNA and RNA/DNA hybrids, an aliquot of the dense DNA fraction of the second gradient was desalted and subsequently digested for 1 h at 37°C with single-strand-specific nuclease P1 and RNAseH (both New England Biolabs). Digestion-resistant DNA was purified by phenol/chloroform extraction and ethanol precipitation, and subsequently separated on a third density gradient. Dense nuclease-resistant DNA fractions were isolated, desalted and subjected to qPCR analysis.

DNA at selected regions of the genome were quantitated in isolated and desalted DNA fractions by quantitative polymerase chain reaction (qPCR) on the iCycler platform using the iTaq SYBR green supermix (BioRad). Primer pairs amplifying replication origins and corresponding adjacent background sites were synthesised by Sigma-Aldrich and used in PCR reactions at 0.5 µM for each primer. The following primer pairs were used:

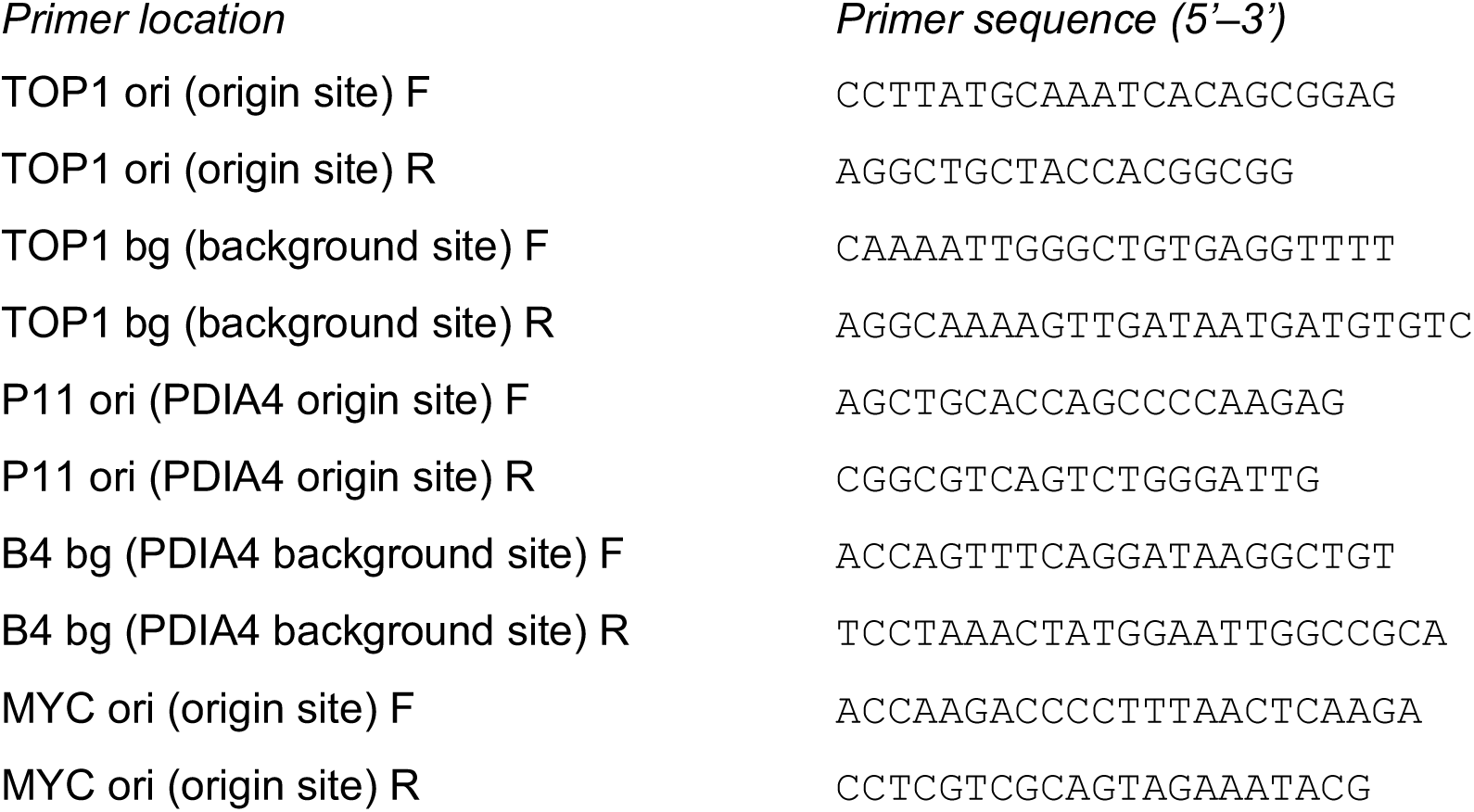

Standard calibration curves for each primer pair were generated with ten-fold serial dilutions of fragmented bulk genomic DNA from quiescent EJ30 cell nuclei. PCR amplification was run after an initial melting at 95°C for 3 min for 42 cycles of 95°C (20 s), 64°C (30 s), 72°C (45 s). DNA amplification was quantitated against the calibration curves. Mean values and standard deviations of at least three reactions were determined.

### PCR-based synthesis of modified DNA

Canonical and modified DNA fragments were synthesised by PCR on 2.5 ng template DNA per reaction. Template DNA was total DNA prepared from quiescent human EJ30 cells fragmented to 250-400 bp. DNA sequences of the synthesised PCR amplicons, as determined by Sanger sequencing, are shown in supplementary Table S1.

Canonical, unmodified DNA fragments were synthesised after an initial melting at 95°C for 3 min for 42 cycles of melting at 95°C (20 s), annealing at 64°C (30 s), and extension at 72°C (45 s), using Taq DNA polymerase in ThermoPol reaction buffer (New England Biolabs), 1 mM MgSO_4_, and 0.2 mM each of dATP, dTTP, dCTP, dGTP (Invitrogen) in a final volume of 50 µl. Fragments were polished by a final extension at 72°C for 7 min. Synthesised DNA fragments were confirmed by agarose gel electrophoresis and Sanger sequencing of both strands, using the same primers as for PCR synthesis (University of Cambridge, Dept. of Biochemistry).

For the synthesis of methylated and further oxidised DNA fragments, the dCTP in the PCR reaction mix was replaced entirely by 0.2 mM of either 5-methyl-dCTP (Jena Bioscience), 5-hydroxymethyl dCTP (Jena Bioscience), 5-formyl dCTP (TriLink Biotechnologies) or 5-carboxy-dCTP (TriLink Biotechnologies). The PCR cycle number and individual step times were as those for canonical DNA synthesis. Synthesis was performed with Deep Vent (exo^-^) DNA polymerase in ThermoPol reaction buffer (New England Biolabs). The melting temperature was raised to 100°C for all DNA fragments. Other PCR parameters were adapted for each fragment individually as follows:

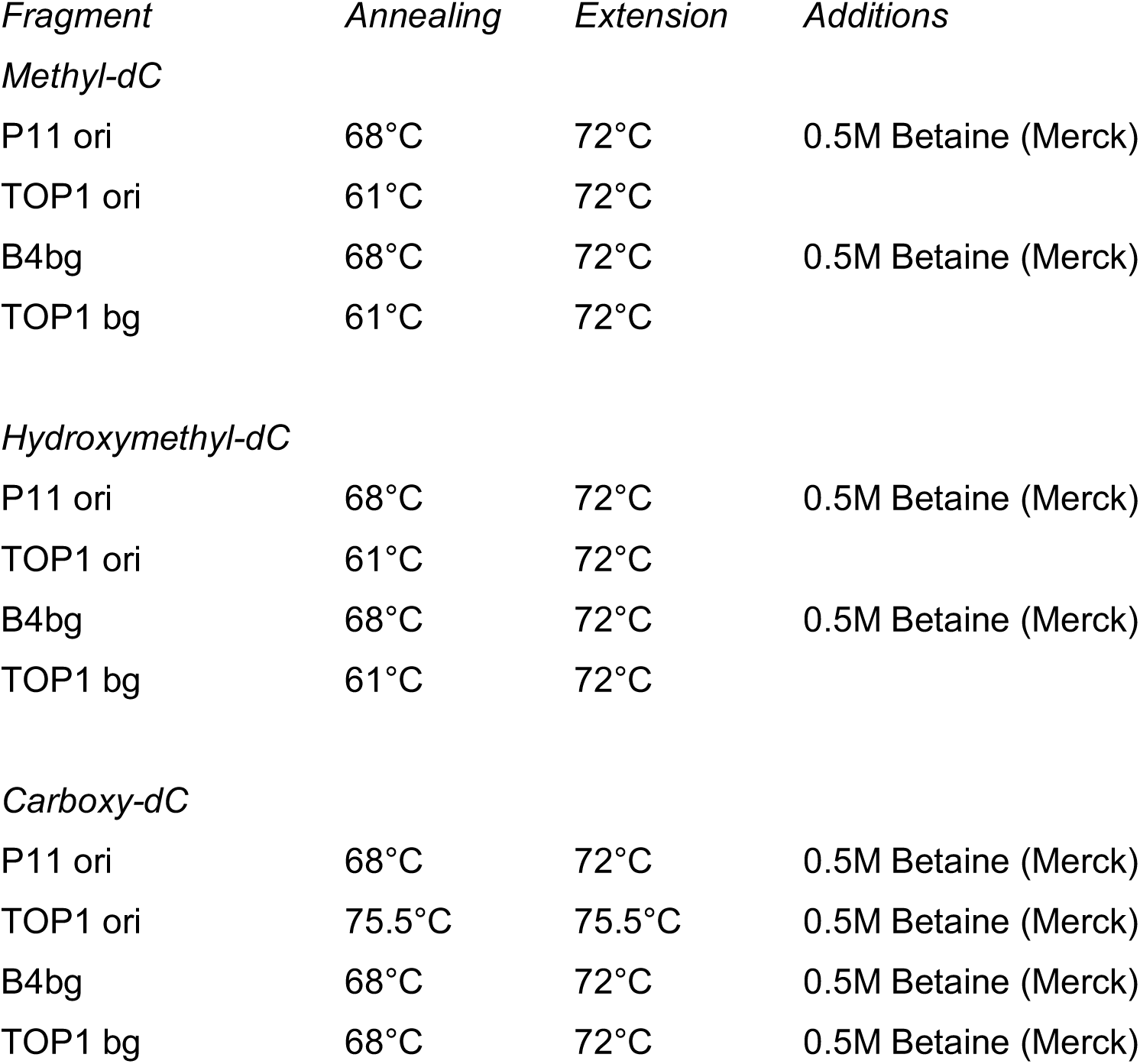

| *Fragment* | *Annealing* | *Extension* | *Additions* |
| --- | --- | --- | --- |
| *Methyl-dC* |  |  |  |
| P11 ori | 68°C | 72°C | 0.5M Betaine (Merck) |
| TOP1 ori | 61°C | 72°C |  |
| B4bg | 68°C | 72°C | 0.5M Betaine (Merck) |
| TOP1 bg | 61°C | 72°C |  |
| *Hydroxymethyl-dC* |  |  |  |
| P11 ori | 68°C | 72°C | 0.5M Betaine (Merck) |
| TOP1 ori | 61°C | 72°C |  |
| B4bg | 68°C | 72°C | 0.5M Betaine (Merck) |
| TOP1 bg | 61°C | 72°C |  |
| *Carboxy-dC* |  |  |  |
| P11 ori | 68°C | 72°C | 0.5M Betaine (Merck) |
| TOP1 ori | 75.5°C | 75.5°C | 0.5M Betaine (Merck) |
| B4bg | 68°C | 72°C | 0.5M Betaine (Merck) |
| TOP1 bg | 68°C | 72°C | 0.5M Betaine (Merck) |

### DNA sequencing

Prior to library generation for Illumina DNA sequencing, isolated and desalted dense and bulk DNA fractions were concentrated by precipitation in 50% isopropanol, 0.5 M NH_4_-acetate, 2.5 µl/ml glycogen (RNA grade, ThermoScientific), washed in 70% ethanol, and dissolved in water.

Libraries for Illumina Sequencing were synthesised using KAPA Hyper kits (Roche) according to the manufacturer’s instructions. High throughput single-end DNA sequencing was performed on an Illumina HiSeq at the Advanced Sequencing Facility, The Francis Crick Institute, London.

Sanger sequencing of PCR products was performed on both DNA strands using the same primers as for PCR, on an Applied Biosystems 3730xl DNA Analyser at the DNA sequencing facility, Department of Biochemistry, University of Cambridge.

### Computational data analysis

Illumina DNA sequencing results were obtained as raw demultiplexed .fastq files after removal of primer sequences and end trimming. Original sequencing results for two independent IniSeq datasets A and B were retrieved as .fastq files (Langley et al., 2016).

Separate .fastq files from one sequencing reaction were first concatenated and then aligned against the human genome hg38 using `bowtie2` (Langmead and Salzberg, 2012). Reads not uniquely mapped were removed and the resulting .sam files were converted to .bam files using `samtools view` (`subcommands: -S -b)` (Li et al., 2009). PCR duplicates were removed by `samtools rmdup` (`subcommand: -s)`, sorted by `samtools sort,` and indexed by `samtools index`. The .bam files were converted to .bed files with `bedtools bamtobed` (Quinlan and Hall, 2010). Read coverage profiles were generated as .bedgraph files from .bed files using `bedtools genomecov (option: -bg)` against a hg38 genome file and then converted to .tdf files with `IGVtools` (Robinson et al., 2011).

Read accumulation peaks were called using .bed files of each experimental sample against the corresponding reference bulk DNA sample file with `MACS2 callpeak` (`subcommands: --nomodel --min-length 200 --max-gap 500 -g 2.7e+9)` (Zhang et al., 2008). Average fragment length values were determined experimentally for each DNA preparation prior to sequencing and implemented into the peak calling via (`--extsize`). A Q-value cutoff (`-q`) of 1.0e-11 was chosen for high stringency.

Intersect analysis of a sample a with a sample b was performed using .bed files with `bedtools intersect (subcommand: -u)`, using intersect distances of 0 bp. Peak positions were randomised for each chromosome with `bedtools shuffle (subcommands: -chrom -noOverlapping)`. Venn diagrams were plotted in R using the `venneuler` package (https://www.R-project.org/).

Coverage profiles, MACS2 peak positions and genomic features were visualised on the `Integrative Genomics Viewer (IGV 2.18)` (Robinson et al., 2011) and further processed as .svg files with `Inkscape 1.1`.

### Mass spectrometry

Nucleosides were prepared by digesting and dephosphorylating up to 1µg of desalted bulk or dense DNA with a Nucleoside Digestion Mix (New England Biolabs) according to the supplier’s specifications in a 50 µl reaction volume containing 1 mM ZnCl_2_, 50 mM Na-acetate, pH 5.4 for 18 h at 37°C. For preparation of nucleosides for calibration, samples containing 1 mM of each dCTP, 5me-dCTP, 5hm-dCTP, 5f-dCTP and 5ca-dCTP were treated identically.

Nucleoside mixtures were analysed by LC-MS (Mass spectrometry service, Yusuf Hamied Department of Chemistry, University of Cambridge), on a Waters VION QTOF mass spectrometer fitted with a Waters UPLC system and a Waters BEH 1.7 µm C18 column at a flow rate of 200 μl/min. For each run, 5 µl of sample was injected in a water:acetonitrile gradient with 0.1% formic acid. The MS spectrum was acquired using an MS^E^ method in positive ionisation mode with capillary voltage set to 0.7 kV, collision energy for the low energy channel was kept at 6 eV and the high energy channel was ramped from 15 to 45 eV. Lock mass correction was applied over the run. Source temperature was set to 120°C and desolvation temperature to 280°C, while cone gas flow was 50 l/h and desolvation gas flow was kept at 800 l/h. Raw spectral data were obtained for material flowing through the column within an elution time of 0-3.5 min. Signal peaks were identified by the Waters UNIFI software and obtained as tables detailing observed mass (m/z) and signal intensity (detector counts) for each detected peak. Nucleoside-specific peaks were identified by analysing calibrator nucleoside mixtures containing combinations of dC, 5me-dC, 5hm-dC, 5f-dC and 5ca-dC and peaks arising from association of nucleosides with H (m/z +1) and Na (m/z +23) were obtained (Fig. S5). In the experimental samples, corresponding nucleoside peaks were identified within a variance of 0.025 Dalton to the calibrator m/z (i.e. < 0.01%). Results are shown as percentages of total deoxycytidines. Signal intensities for H and Na associated peaks were combined for each nucleoside. Mean values were obtained for n=2-5 repeats.

### Detection of 5’-hydroxymethylcytosine by MspI-sensitivity

Purified and desalted bulk and dense DNA fractions were first glucosylated at hydoxymethylcytosine residues before subjected to restriction endonuclease digestion of non-glucosylated DNA. The DNA (25 – 100 ng) was incubated in a reaction volume of 50 µl at 37°C for 18 h with 10 U of T4 beta-glucosyltransferase (T4-BGT) in the presence of 80 µM UDP-glucose in NEBuffer 4 (all New England Biolabs). Control reactions were incubated without T4 beta-glucosyltransferase. These DNA samples were then digested at 37°C for 6 h with 100 U MspI (New England Biolabs). Reactions were inactivated with Proteinase K at 42°C for 45 min, and then by incubation at 95°C for 15 min. The presence of uncut template DNA was determined by PCR.

## Supporting information

Supplementary figures, table and their legends

## Acknowledgements

We thank Ron Laskey, Guillaume Guilbaud, Pierre Murat, and Clara Collart for discussions and for critical reading of the manuscript.

We thank E. Roberto Canales Candela and Dijana Matak Vinkovic at the Yusuf Hamied Department of Chemistry, University of Cambridge, for discussions and mass spectrometry services.

This work was supported by core and teaching funds of the Department of Zoology, University of Cambridge. J.B. and R.D. were furthermore supported by undergraduate student summer scholarships from Christ’s College, Cambridge.

We were also funded by the Francis Crick Institute, which receives its core funding from Cancer Research UK (FC001157), the UK Medical Research Council (FC001157), and the Wellcome Trust (FC001157).

We are grateful to the Francis Crick Institute Advanced Sequencing facility, which receives funding from Cancer Research UK (CC1064), the UK Medical Research Council (CC1064), and the Wellcome Trust (CC1064).

The work benefited from the Imaging Facility, Department of Zoology, supported by funds from a Wellcome Trust Equipment Grant (WT079204) with contributions by the Sir Isaac Newton Trust in Cambridge, including Research Grant [18.07ii(c)].

## Author contributions

Conceptualisation, T.K.; Methodology, T.K.; Investigation, T.K., J.B., R.D, R.A.J.; Writing – Original Draft, T.K.; Writing – Review & Editing, T.K. and J.C.S.; Visualisation, T.K.; Supervision, T.K. and J.C.S.; Funding Acquisition, T.K. and J.C.S.

## Data Availability

Raw and processed DNA sequencing data will be shared after peer review.

