## Supplementary figures, table and their legends for "Human DNA replication initiation sites are specified epigenetically by oxidation of 5-methyl-deoxycytidine"

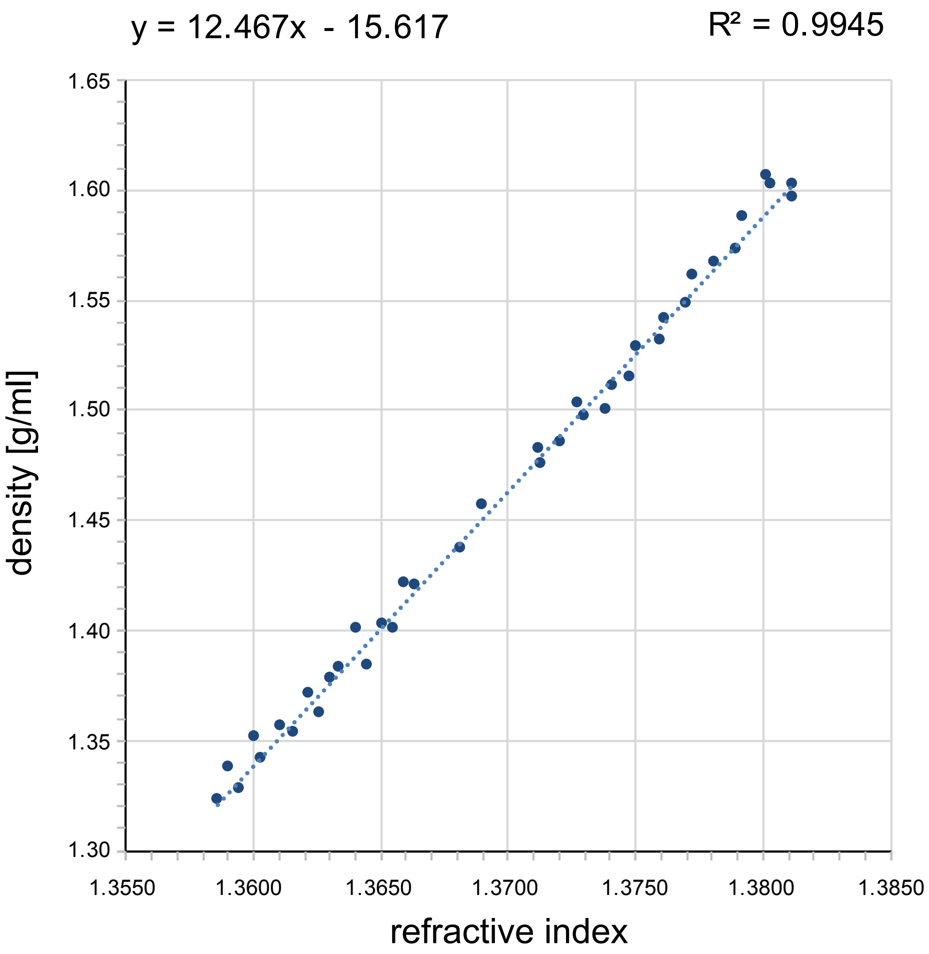


**Figure S1**

**Conversion of refractive index to density**

Plot of experimentally determined densities (D) and refractive indexes (RI) of n=40 fractions across two caesium sulphate density equilibrium gradients. The linear regression follows the equation: D = 12.46 x RI x g/ml -15.62 ($R^2$ = 0.9945).

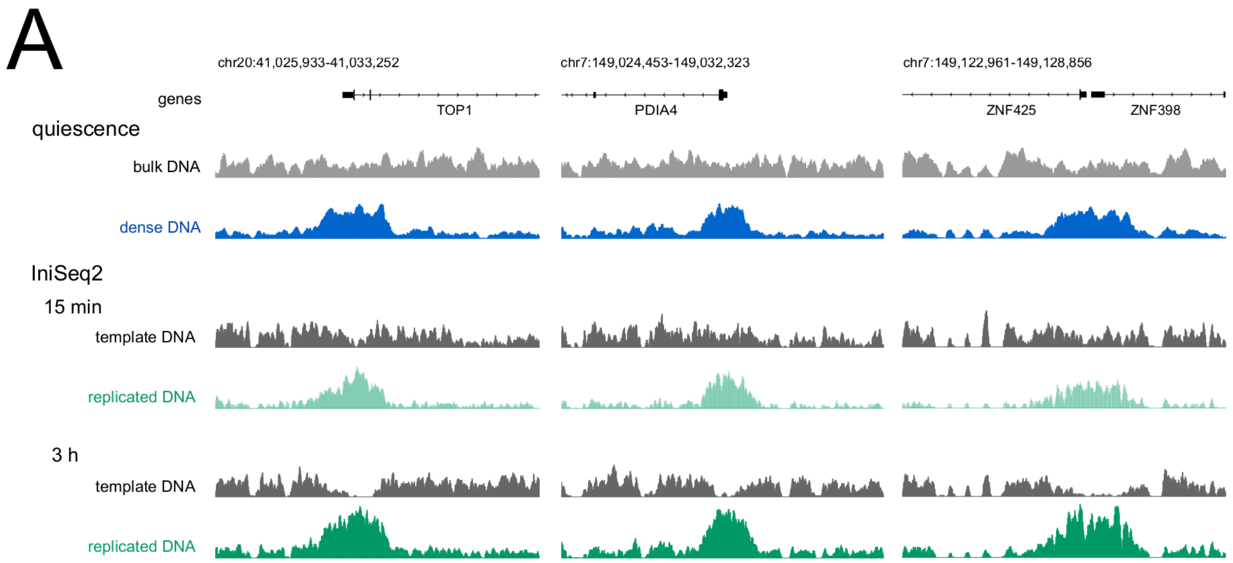


**Figure S2**

**Naturally dense DNA is enriched at active and efficient DNA replication origins**

Integration of dense DNA with original IniSeq2 data (Guilbaud et al., 2022). **(A)** Illumina sequencing read coverage profiles of bulk and dense DNA from quiescent cells (grey and blue) are compared to IniSeq2 profiles of unreplicated template DNA (grey) and replicated DNA (green) at the TOP1 (left), PDIA4 (middle) and ZNF425/398 origin sites (right) after 15 min and 3 h incubations. Genome coordinates and positions of reference genes are indicated. Note the local reduction of unreplicated template DNA and concomitant accumulation of replicated DNA over replication time at origin sites marked in quiescence by dense DNA.

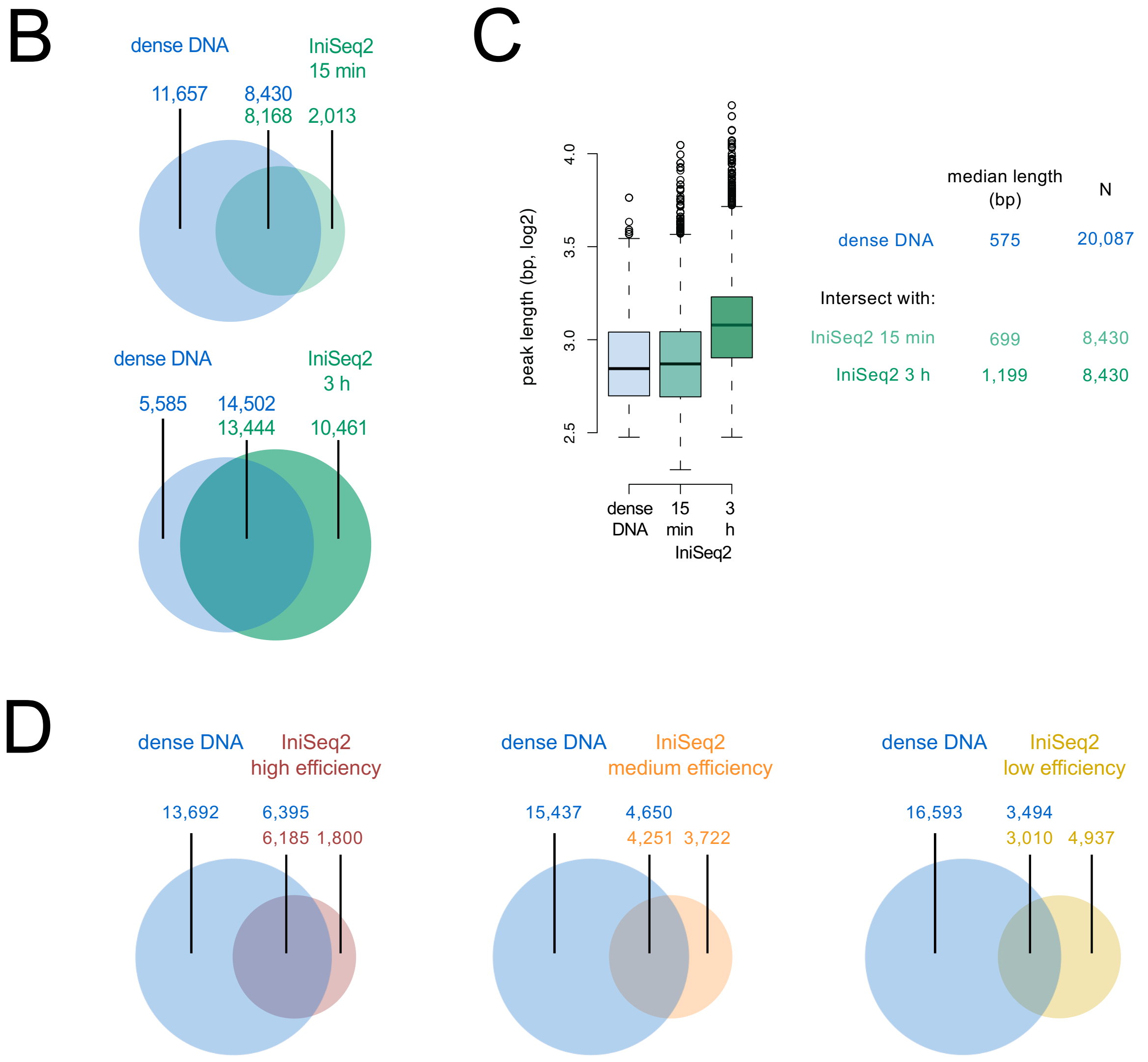


**Figure S2, continued**

**Naturally dense DNA is enriched at active and efficient DNA replication origins 2**

**(B)** Genome-wide intersect analyses between MACS2 peaks of dense DNA and Ini-Seq2-seq DNA replication origin sites determined by a custom algorithm based on the conversion of unreplicated DNA to replicated DNA after 15 min and 3 h incubation time (Guilbaud et al., 2022). Non-intersected and intersected numbers of peaks are indicated; the top number in the intersect indicates the number of peaks of the left distribution (dense DNA) intersecting with the right distribution (IniSeq2 origins), and the bottom number indicates the number of peaks of the right intersecting with the left distribution. **(C)** Distributions of peak lengths for dense DNA in quiescent cells (blue) intersecting with IniSeq2 origins after 15 min (pale green), and again at 3 h incubation time (green). Tabulated median and N values for these distributions are included on the right. **(D)** Intersect analyses between MACS2 peaks of dense DNA and high-, medium- and low-efficiency class IniSeq2 origins (Guilbaud et al., 2022).

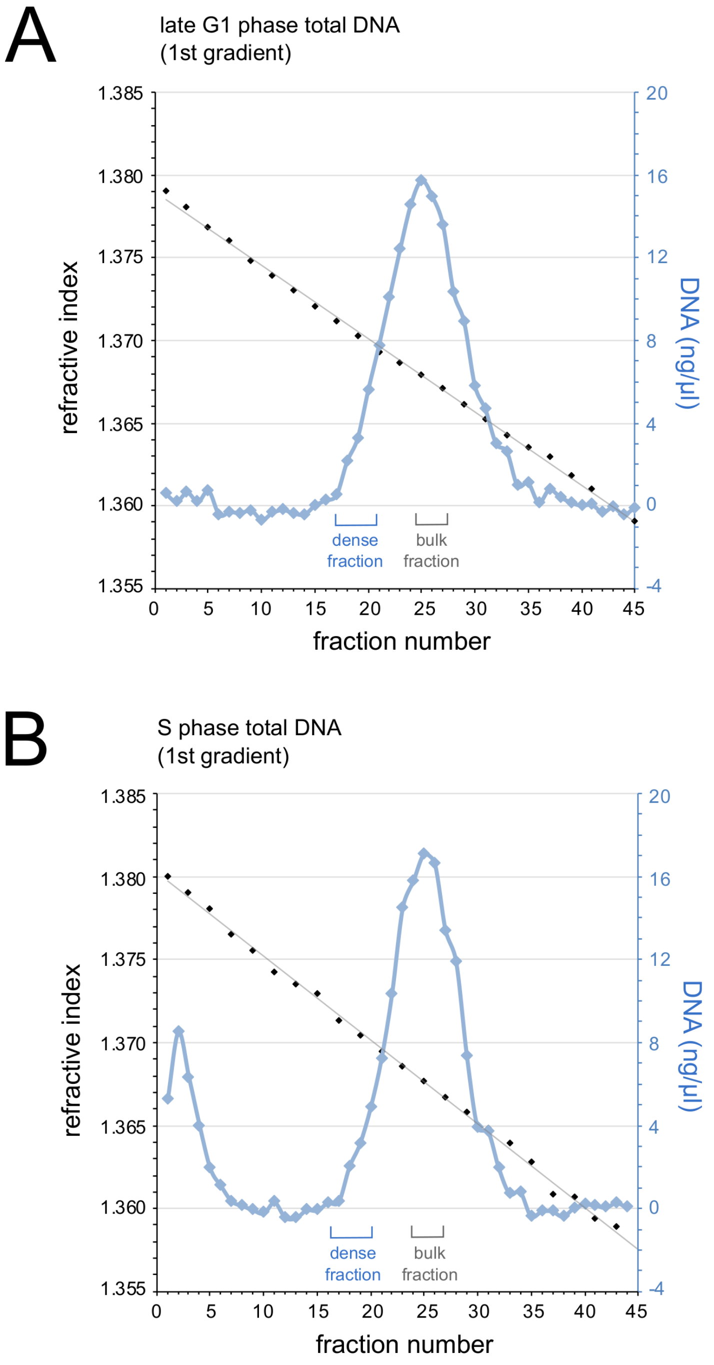


**Figure S3**

**Density gradient separation of fragmented late G1 and S phase human DNA**

Human EJ30 cells were synchronised in late G1 phase by mimosine (Krude, 1999) and in early S phase by thymidine (Krude et al., 1997). Genomic DNA was purified, fragmented and separated on primary caesium sulphate density equilibrium gradients. **(A)** Separation of late G1 phase cell DNA. Isolated dense DNA fractions for loading onto a second gradient and isolated bulk DNA fractions selected for sequencing and PCR analysis are indicated. **(B)** Separation of S phase cell DNA. See Figure 3 for secondary density gradients and further analysis.

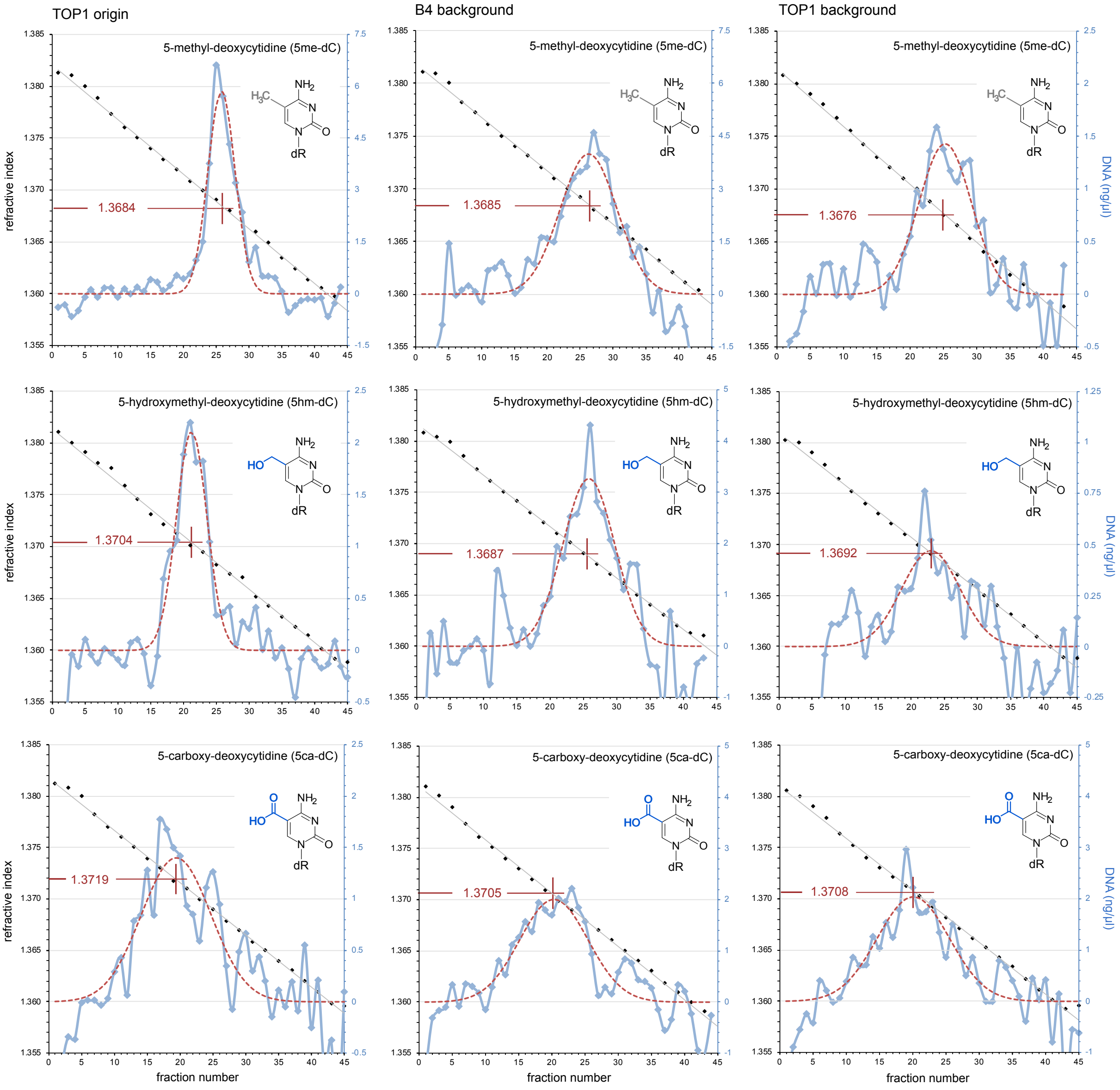


### Figure S4

### Dense DNA is a consequence of oxidised methyl-deoxycytidines

Comparative density analysis of methylated and further oxidised DNA fragments of the TOP1 origin and B4 and TOP1 background sites. PCR products were synthesised using modified dCTP for an incorporation of 5me-dC (top row), 5hm-dC (middle row), and 5ca-dC (bottom row). DNA fragment identities, deoxycytidine modifications, DNA concentrations (pale blue), together with fitted Gauss distributions (hatched red lines), and the refractive indexes for the means of each fitted distribution are indicated. Note the increased densities for DNA fragments containing 5hm-dC and 5ca-dC. See Table S1 for DNA sequences of these fragments.

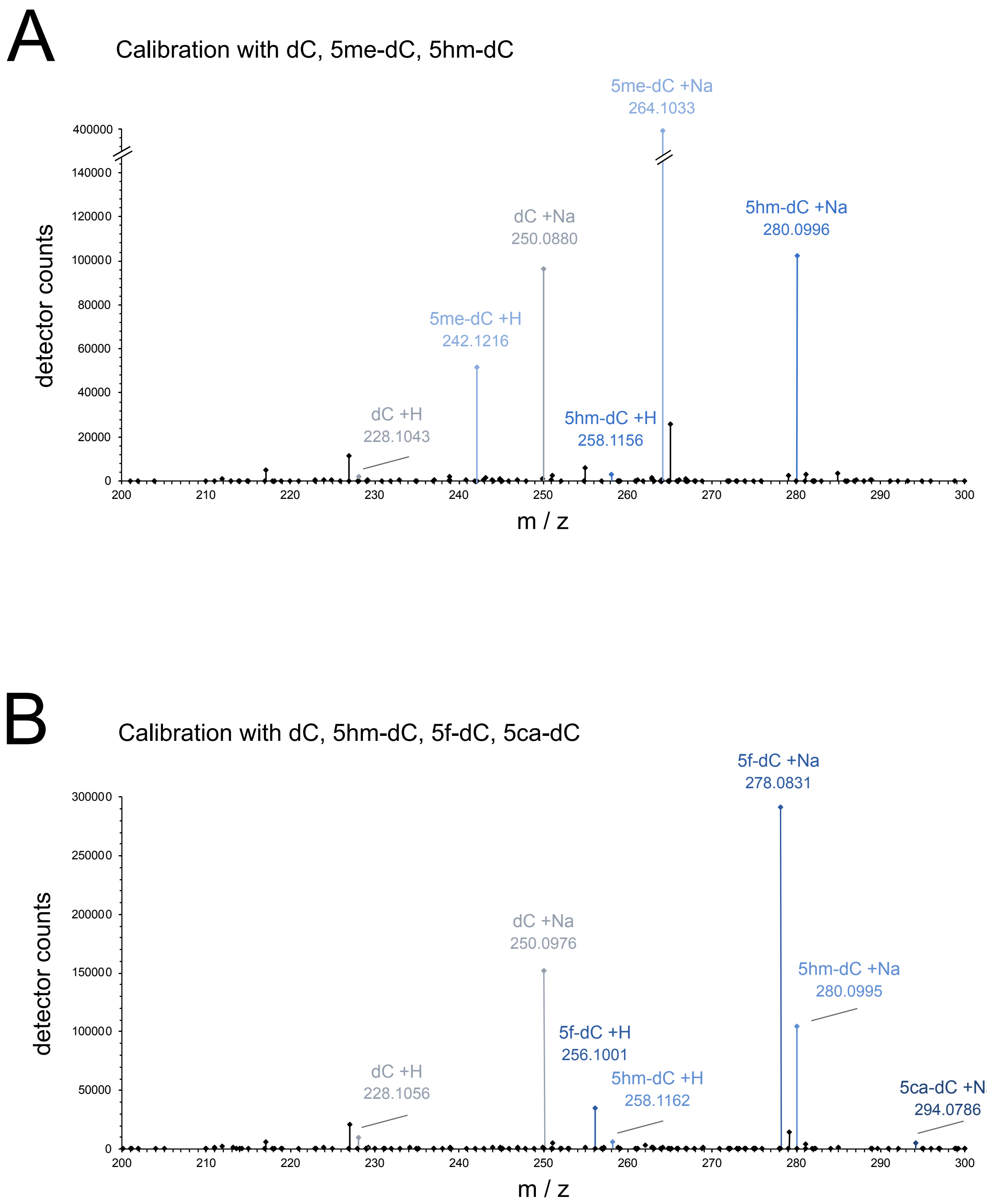


**Figure S5**

**Calibration of mass spectra using defined deoxycytidine derivatives**

Two separate combinations of nucleoside triphosphates were dephosphorylated and subjected to LS-MS: **(A)** dCTP, 5me-dCTP, 5hm-dCTP. **(B)** dCTP, 5hm-dCTP, 5f-dCTP and 5ca-dCTP. Plots of detector counts across the mass-over-charge (m/z) range of 200-300 Da are shown. Peaks corresponding to the monoisotopic masses of the indicated nucleosides including H and Na adducts are highlighted.

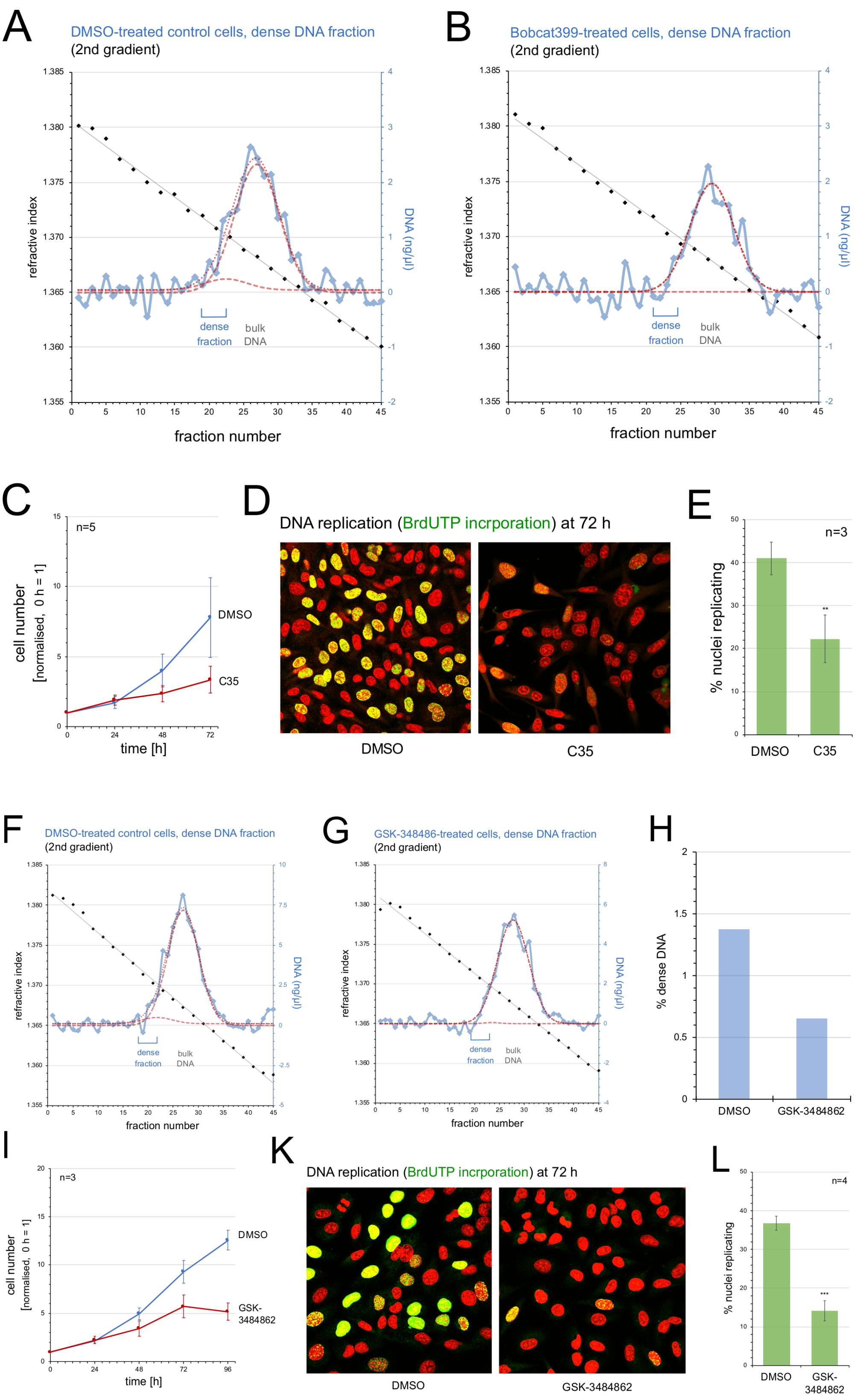


**Figure S6**

**Methylation of deoxycytidine and its further oxidation is required for cell proliferation and DNA replication**

**(A, B)** Density gradient analysis of fragmented DNA from DMSO and Bobcat399-treated cells.

**(A)** Separation of dense DNA from asynchronously proliferating, DMSO-treated cells on a second caesium sulphate density gradient. **(B)** Separation of dense DNA from cells treated with 125 µM of Bobcat399 on a second caesium sulphate density gradient. Calculated best fits to two normal Gaussian distributions are plotted by hashed (individual) and dotted (combined) red lines for dense and bulk DNA. Positions of reference bulk DNA, and of isolated dense DNA fractions selected for further analysis (shown in Fig. 6B) are indicated.

**(C-E)** Inhibition of TET enzyme activity by C35. **(C)** Inhibition of cell proliferation. Asynchronously proliferating EJ30 cells were treated with DMSO (blue) or 150 µM of C35 (red) for 72 h. Normalised cell numbers are plotted against time as averages ± standard errors of the mean for n=5 independent experiments. **(D, E)** Inhibition of DNA replication. After 72 h of treatments, replicating cells were pulse-labelled with BrdU and replicating cell nuclei detected by confocal immunofluorescence microscopy. **(D)** Representative micrographs. DNA is stained with propidium iodide (red), BrdU incorporation with specific antibodies (green). Note the lower proportions of replicating cell nuclei and lower intensity of BrdU incorporation signals in C35-treated cells compared to the DMSO control. **(E)** Quantification of the percentages of replicating cell nuclei. Proportions of nuclei replicating their DNA were scored and plotted. Mean values ± standard errors of the mean from n=3 independent preparations are shown (T-tests, two-tailed, unequal variance between DMSO control and treatment: **, $p<0.01$).

**(F-L)** Inhibition of DNMT activity by GSK3484862. **(F)** Separation of dense DNA from asynchronously proliferating, DMSO-treated cells on a second caesium sulphate density gradient. Calculated best fits to two normal Gaussian distributions are plotted by hashed (individual) and dotted (combined) red lines for dense and bulk DNA, and isolated dense DNA fractions selected for further analysis are indicated. **(G)** Separation of dense DNA from cells treated with 10 µM of GSK3484862 on a second caesium sulphate density gradient. **(H)** Quantification of relative dense DNA amounts. Relative amounts were obtained as percentages of dense DNA out of total DNA from the DMSO and GSK3484862-treated cells. **(I)** Inhibition of cell proliferation. Asynchronously proliferating EJ30 cells were treated with DMSO (blue) or 10 µM of GSK3484862 (red) for 96 h. Normalised cell numbers are plotted against time as averages ± standard errors of the mean for n=3 independent experiments. **(K, L)** Inhibition of DNA replication. After 72 h of treatment, replicating cells were pulse-labelled with BrdU and replicating cell nuclei detected by confocal immunofluorescence microscopy. **(K)** Representative micrographs. Note the lower proportions of replicating cell nuclei and lower intensity of BrdU incorporation signal in GSK3484862-treated cells compared to the DMSO control. **(L)** Quantification of the percentages of replicating cell nuclei. Proportions of nuclei replicating their DNA were scored and plotted. Mean values ± standard errors of the mean from n=4 independent preparations are shown (T-tests, two-tailed, unequal variance between DMSO control and treatment: ***, $p<0.001$).

**Table S1**

DNA sequences of amplified PCR products (5’ – 3’).
Additional Adenosine residues added to the 3’ end of each strand during PCR by Taq polymerases are shown in grey. MspI restriction sites are indicated in bold.

TOP1 ori (TOP1 origin site)

TCCTTATGCAAATCACAGCGGAGCGCGCACGGT**CCGG**AGGCGGGGCTTGCGATGCAAAGACA
GGTCCGTCTGGCGAACAGCGAGGGGGCGGGCCGCAACCCTCTGCCTCTTTCCGCGAGCGCTG
ACGTCGCCGACGTGTTGTTTAAAAGCGGCCGCGCAGGCGCAGTGAGCCCAAATGCGAACTTA
GGCTGTTACACAACTGCTGGGGTCTGTTCTCGCCGCCCGC**CCGG**CAGTCAGGCAGCGTCGCC
GCCGTGGTAGCAGCCTA

(265bp, 64%GC)

TOP1 bg (TOP1 background site)

TCAAAATTGGGCTGTGAGGTTTTTTTGTTTTGCTTGTTTTCAGTGCATGTGTGGGGTGGGGA
GGTGCAAAAAAATGTTTGCCTTCTAATATACAGCTTTGTTGTATATTAATTATATAGTAACA
GTTGTCCCTGCTGCTAATGGTATGGAAAATGATTCTGTTGTGCTTTCTGAATACATATCATT
TTAGAAGTTTCAGATTATACTAAAACCATGTTATTTCATATTAGTTGGACATGAATGCATAT
TTATATGACACATCATTATCAACTTTTGCCTA

(280bp, 31%GC)

P11 ori (PDIA4 origin site)

TAGCTGCACCAGCCCCAAGAGCAGCAGGAGCAGGAAGGCTTT**CCGG**GGCCTCATGGTAGCGG
GGGCGGAGCGCGGCCTCCTAGCGTCGGCGGCCGCTGAGCGCACCGAGAACTCGGGGTCTGGC
CGACAGCCCGTCGCTCCTTAGCGACGCGGGGGAG**CCGG**AAAAACCCACGGAAGTCGTCCCCG
GCGATTGGCAGGGGGCGGAGGAAGTCGCGGG**CCGG**CCAATCCCAGACTGACGCCGA

(242bp, 70%GC)

B4 bg (PDIA4 background site)

TACCAGTTTCAGGATAAGGCTGTTTCTTTTTTTTAGATATTTGCTGAGGGCTGAAACAAGCA
AGCTGTTAATGCCTGAACTTTTTTCCTACCAAGTCCCTAGTGCTTCATATTAGTATGTACAT
GGTCAGGTCCCAAGAAACAGCAGGTGATAGCTCCCACCCAACAAAGACTGTCAACTTCCTGA
ACCAGGACAGTCCTCACTTCCCTCCATCCCACTCTTTGCGGCCAATTCCATAGTTTAGGAA

(247bp, 43%GC)
